# Differences in the inflammatory proteome of East African and Western European adults and associations with environmental and dietary factors

**DOI:** 10.1101/2022.08.23.504992

**Authors:** Godfrey S. Temba, Nadira Vadaq, Vesla Kullaya, Tal Pecht, Paolo Lionetti, Duccio Cavalieri, Joachim L. Schultze, Reginald Kavishe, Leo A.B. Joosten, Andre J. van der Ven, Blandina T. Mmbaga, Mihai G. Netea, Quirijn de Mast

## Abstract

Non-communicable diseases (NCDs) are rising rapidly in urbanizing populations in sub-Saharan Africa. Assessment of inflammatory and metabolic characterstics of an urbanizing African population and the comparison with populations outside Africa could provide insight in the pathophysiology of the rapidly increasing epidemic of NCDs, including the role of environmental and dietary changes. Using a proteomic plasma profiling approach comprising 92 inflammation-related molecules, we examined differences in the inflammatory proteome in healthy Tanzanian and healthy Dutch adults. We show that healthy Tanzanians display a pro-inflammatory phenotype compared to Dutch subjects, with enhanced activity of the Wnt/β-catenin signalling pathway and higher concentrations of different metabolic regulators such as 4E-BP1 and fibroblast growth factor 21. Among the Tanzanian volunteers, food-derived metabolites were identified as an important driver of variation in inflammation-related molecules, emphasizing the potential importance of lifestyle changes. These findings endorse the importance of the current dietary transition in the NCDs epidemic in sub-Saharan Africa and the inclusion of underrepresented populations in systems immunology studies.

## Introduction

The human immune response is tightly regulated by a complex and intricate network of pro-and anti-inflammatory cytokines, chemokines and other immuno-metabolic mediators. Prolonged dysregulation of this network may result in an unresolved pro-inflammatory state, which is central to the development of a wide range of non-communicable diseases (NCDs), including cardiovascular disease, diabetes, rheumatic and inflammatory bowel diseases, and even malignancies (Choi et al., 2015; Moore & Tabas, 2011; Seyedsadjadi & Grant, 2021). The inflammatory response is highly variable among healthy individuals as a consequence of genetic and non-genetic factors (Brodin & Davis, 2017); this includes intrinsic factors such as age and sex, as well as environmental exposures such as diet and past infections (Liston, Humblet-Baron, Duffy, & Goris, 2021). Not surprisingly, variation in inflammatory proteins between populations has also been reported (A. Schutte, 2006; A. E. Schutte, Myburgh, Olsen, Eugen-Olsen, & Schutte, 2012). Understanding the nature of this variation and the factors involved is key for understanding the dynamics of infectious as well as immune-mediated pathology across populations.

Worldwide, many communities are currently undergoing a rapid urbanization process with a transition of lifestyle, and diet, characterized by a more sedentary lifestyle and a shift from traditional high-fiber diets to a diet richer in processed foods, animal fat and simple carbohydrates. Also, environmental exposures change from close contact with animals and smoke from burning wood to petrol gasses. This is accompanied by an epidemiologic transition in which the burden of disease shifts from infectious diseases to NCDs (Beaglehole et al., 2011; Unwin et al., 2010). This epidemiologic transition is at least partly mediated through effects on the immune system of individuals across communities. We recently reported that urban-living Tanzanians display a pro-inflammatory gene signature and higher *ex-vivo* cytokine responses compared to rural-living individuals (Temba et al., 2021) and that food-derived metabolites are an important driver of this difference. In communities in sub-Saharan Africa, this effect may also be more pronounced than in other populations, as the historically high burden of infectious diseases may have resulted in the selection of genotypes favoring a robust immune response (Karlsson, Kwiatkowski, & Sabeti, 2014). Indeed, we recently showed important differences in the genetic regulation of cytokine responses between healthy Tanzanian and Dutch individuals, with enrichment of interferon pathways in the Tanzanians (Boahen et al., 2022). In a context with a declining burden of infectious diseases, alongside a shift to an unhealthy Western-type lifestyle, such a heritable pro-inflammatory phenotype may particularly drive a health-to-disease transition with the onset of NCDs (Bickler et al., 2018).

Studies on non-Western populations outside historically wealthy countries are underrepresented in systems-immunology literature, despite the fact that these populations are representative of the majority of the world’s population. In addition, these populations offer unique opportunities to increase our understanding of the pathophysiology of NCDs, including the role of diet and environmental exposures. Our present knowledge of the regulation of the immune system in individuals in sub-Saharan Africa and how this compares to individuals in the industrialized world is limited. We investigated the hypothesis that healthy individuals in East Africa have a pro-inflammatory phenotype in comparison to individuals from North-western Europe and that these differences are partly driven by common environmental factors, including diet. Using a 92-plex proteomic panel based on a proximity extension assay technology, we compared the inflammatory proteome of healthy Tanzanians of African origin residing in Tanzania with that of healthy individuals of Western-European ancestry living in the Netherlands. Next, we studied associations between the proteome with intrinsic factors and environmental exposures, with special emphasis on the food-derived metabolites.

## RESULTS

### Demographics

Data from plasma samples of 318 Tanzanian and 416 Dutch individuals were included in this study. Characteristics of study participants are summarized in Table 1 **&** Supplemental Fig. 1. The Tanzanians had a significantly higher median (IQR) age (30.2 years; 23.4-39.9) than the Dutch (23.0 years; 21.0-26.0; *P*-value <0.0001) (Supplemental Fig. 1a), although both populations were young adults. Tanzanian females also had a higher BMI than Dutch females (25.7; 22.6-29.9 vs. 21.5; 20.4-23.1; *P*-value <0.0001).

**Table 1.**
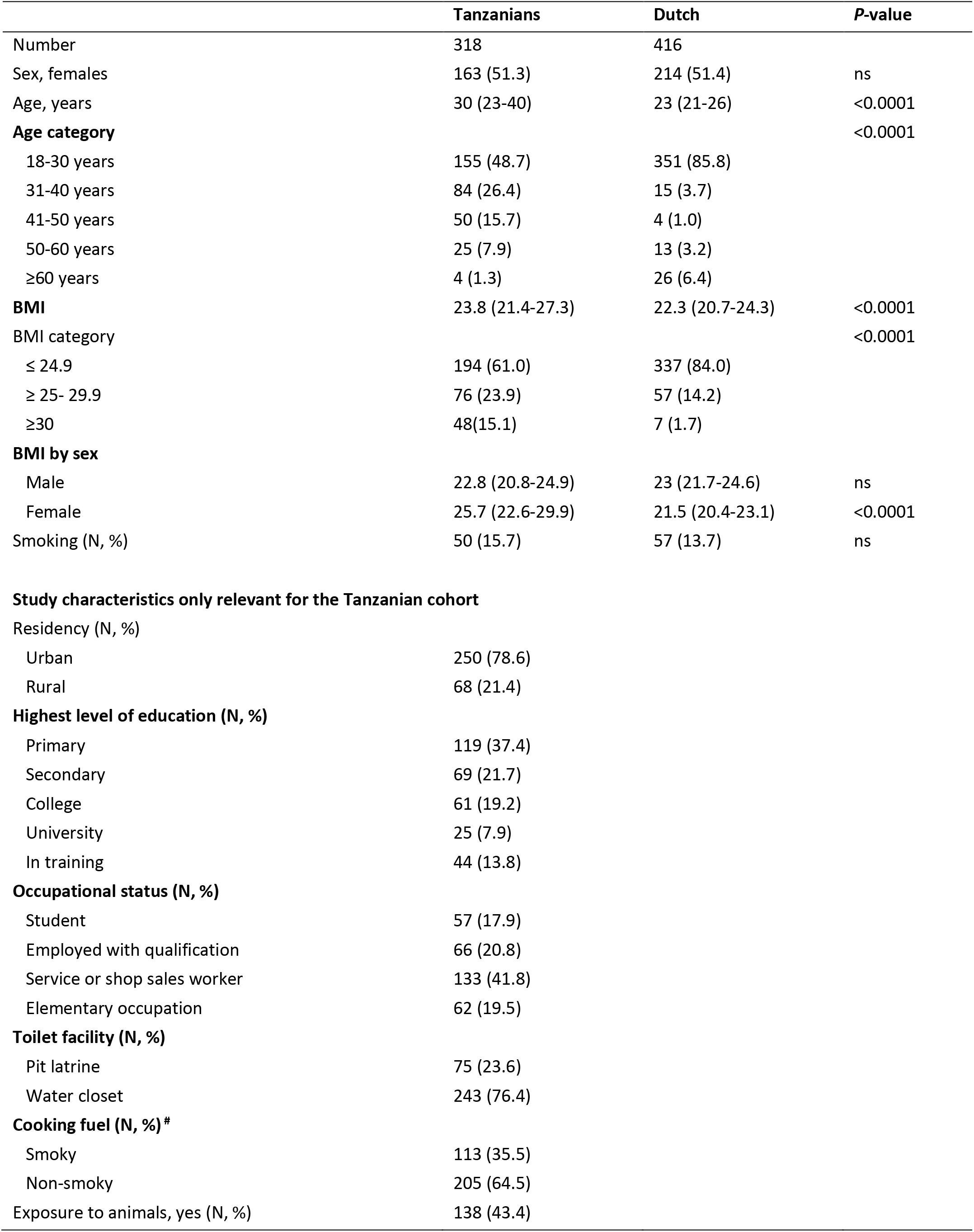
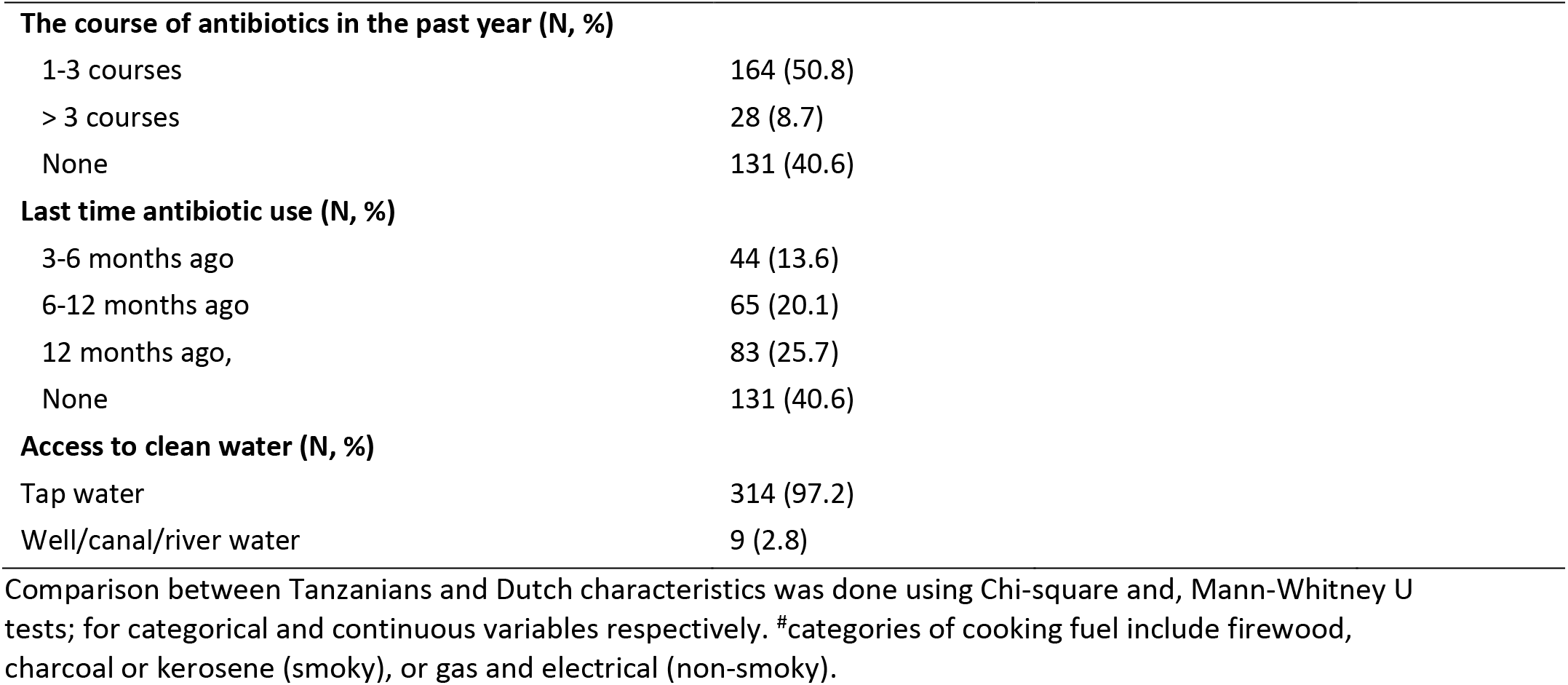
Descriptive characteristics of study participants

|  | **Tanzanians** | **Dutch** | **P-value** |
| --- | --- | --- | --- |
| Number | 318 | 416 |  |
| Sex, females | 163 (51.3) | 214 (51.4) | ns |
| Age, years | 30 (23-40) | 23 (21-26) | <0.0001 |
| **Age category** |  |  | <0.0001 |
| 18-30 years | 155 (48.7) | 351 (85.8) |  |
| 31-40 years | 84 (26.4) | 15 (3.7) |  |
| 41-50 years | 50 (15.7) | 4 (1.0) |  |
| 50-60 years | 25 (7.9) | 13 (3.2) |  |
| ≥60 years | 4 (1.3) | 26 (6.4) |  |
| **BMI** | 23.8 (21.4-27.3) | 22.3 (20.7-24.3) | <0.0001 |
| **BMI category** |  |  | <0.0001 |
| ≤ 24.9 | 194 (61.0) | 337 (84.0) |  |
| ≥ 25- 29.9 | 76 (23.9) | 57 (14.2) |  |
| ≥30 | 48(15.1) | 7 (1.7) |  |
| **BMI by sex** |  |  |  |
| Male | 22.8 (20.8-24.9) | 23 (21.7-24.6) | ns |
| Female | 25.7 (22.6-29.9) | 21.5 (20.4-23.1) | <0.0001 |
| Smoking (N, %) | 50 (15.7) | 57 (13.7) | ns |
| **Study characteristics only relevant for the Tanzanian cohort** |  |  |  |
| **Residency (N, %)** |  |  |  |
| Urban | 250 (78.6) |  |  |
| Rural | 68 (21.4) |  |  |
| **Highest level of education (N, %)** |  |  |  |
| Primary | 119 (37.4) |  |  |
| Secondary | 69 (21.7) |  |  |
| College | 61 (19.2) |  |  |
| University | 25 (7.9) |  |  |
| In training | 44 (13.8) |  |  |
| **Occupational status (N, %)** |  |  |  |
| Student | 57 (17.9) |  |  |
| Employed with qualification | 66 (20.8) |  |  |
| Service or shop sales worker | 133 (41.8) |  |  |
| Elementary occupation | 62 (19.5) |  |  |
| **Toilet facility (N, %)** |  |  |  |
| Pit latrine | 75 (23.6) |  |  |
| Water closet | 243 (76.4) |  |  |
| **Cooking fuel (N, %) #** |  |  |  |
| Smoky | 113 (35.5) |  |  |
| Non-smoky | 205 (64.5) |  |  |
| Exposure to animals, yes (N, %) | 138 (43.4) |  |  |

|  |  |
| --- | --- |
| **The course of antibiotics in the past year (N, %)** |  |
| 1-3 courses | 164 (50.8) |
| > 3 courses | 28 (8.7) |
| None | 131 (40.6) |
| **Last time antibiotic use (N, %)** |  |
| 3-6 months ago | 44 (13.6) |
| 6-12 months ago | 65 (20.1) |
| 12 months ago, | 83 (25.7) |
| None | 131 (40.6) |
| **Access to clean water (N, %)** |  |
| Tap water | 314 (97.2) |
| Well/canal/river water | 9 (2.8) |

Comparison between Tanzanians and Dutch characteristics was done using Chi-square and, Mann-Whitney U tests; for categorical and continuous variables respectively. #categories of cooking fuel include firewood, charcoal or kerosene (smoky), or gas and electrical (non-smoky).

### Differences in inflammatory proteome between Tanzanians and Dutch

The inflammatory proteome was measured simultaneously in samples from the Tanzanian and Dutch participants using the Olink ‘inflammation’ panel, which targets 92 cytokines, chemokines and other inflammation and metabolism-related proteins (Olink® Proteomics AB, Uppsala, Sweden). Relative protein concentrations are reported as normalized protein expression (NPX) units, which are on a Log2 scale. Eighteen proteins were excluded from further analysis because their value was below the lower limit of detection in more than 25% of samples in both cohorts (Supplemental Fig. 2). Principal component analysis (PCA) of the remaining 74 proteins revealed a clear separation between Tanzanian and Dutch samples (Fig. 1a). A volcano plot of differentially expressed proteins (Fig. 1b) showed that 35 (47%) proteins were significantly higher in the Tanzanians and 20 (27%) lower at an FDR *P*<0.05, with correction for age and sex and BMI. The most prominently (fold change (FC)) upregulated proteins were two regulators of metabolism: the mTOR substrate and translational repressor 4E-BP1 (log_2_ FC 1.9; FDR *P*=1.3 x 10^-60^) and FGF21 (fibroblast growth factor 21; log_2_ FC 1.3; *P*=1.3 x 10^-30^), a hormone produced by the liver that functions as a major regulator of glucose and lipid homeostasis. Obesity and excess carbohydrate and/or insufficient protein intake were reported to increase FGF21 concentrations (Hill, Berthoud, Münzberg, & Morrison, 2018). Other prominently upregulated proteins in the Tanzanians were interleukin (IL)-17A (log_2_ FC 0.7; *P*=9.6 x 10^-34^) and IL-17C (log_2_ FC 0.7; *P*=1.1 x 10^-74^), and the CC-chemokine family members CCL11/eotaxin (eosinophil chemoattractant; log_2_ FC 0.7; *P*=7.3 x 10^-63^), CCL3/MIP-1α (macrophage inflammatory protein-1α; log_2_ FC 0.6; *P*=5.6 x 10^-10^), CCL7/MCP3 (monocyte chemotactic protein 3; log_2_ FC 0.6; *P*=7.4 x 10^-42^) and CCL8/MCP2 (log_2_ FC 0.5; *P*=8.4 x 10^-23^). The cytokines Tumour Necrosis Factor (TNF), IL-6, IL-10, and IL-18, as well as oncostatin-M (OSM) and adenosine deaminase (ADA) were also significantly higher in the Tanzanians.

**Figure 1.**
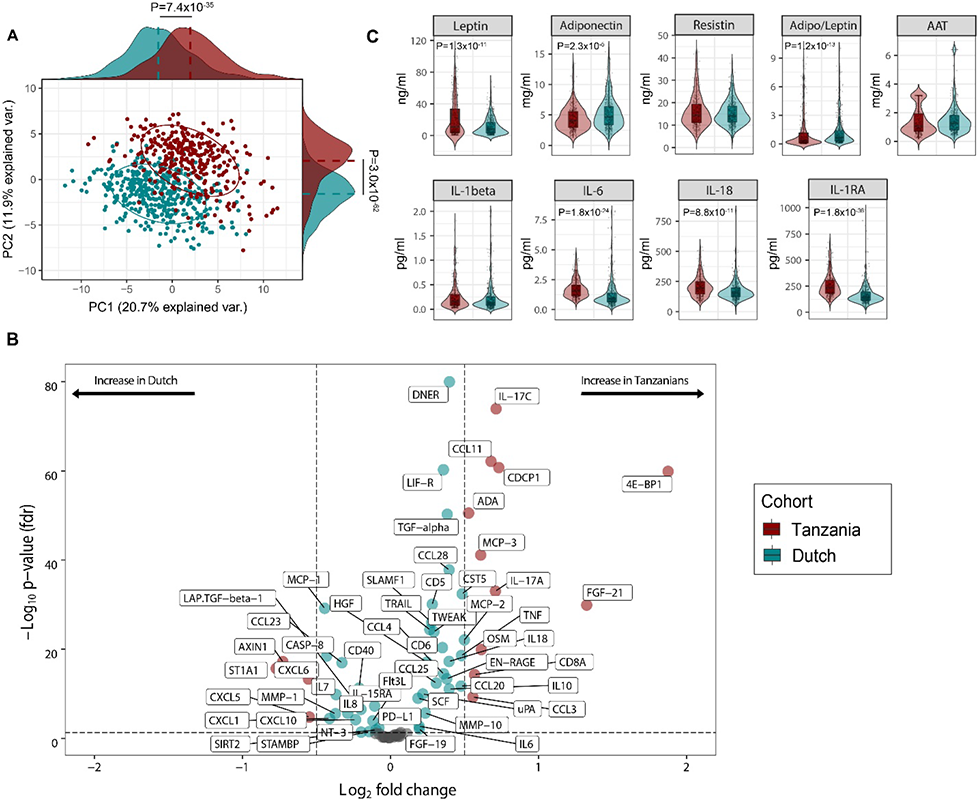
Differentially expressed inflammatory protein profiles among Dutch and Tanzanian participants. **(A)** Principal component analysis depicting the sample distribution of Dutch (N=416) vs. Tanzanian (N=318) healthy individuals across PC1 and PC2, **(B)** Volcano plot of the Dutch and Tanzanian cohorts showing differentially expressed protein (DEPs) analyzed by Limma (linear models for microarray data) R package (Dutch cohort; N=74 and Tanzanian cohort N=72 inflammatory proteins). Presented in the x-axis is the Log_2_ fold change (Log2 FC) of the normalized protein expression (NPX) while the y-axis presents the −Log_10_ of the adjusted p-values (FDR <0.05), dotted lines represent the cut-off value Log2FC <.5 and >.5 **(C)** Violin plots showing the mean (linear regression corrected for age, sex and BMI) differences of the circulating inflammatory cytokines and adipokines in the Dutch and Tanzanian participants (also previously reported (Temba et al., 2022)). Dark red and turquoise colors indicate higher concentrations in the Tanzanian vs. Dutch participants respectively. Results were declared significant after correcting for multiple testing using False discovered rate (FDR); p-value <0.05(*), <0.005(**), and <0.0001(***). Abbreviation: AAT; alpha-1 antitrypsin; BMI; Body Mass Index

**Figure 2:**
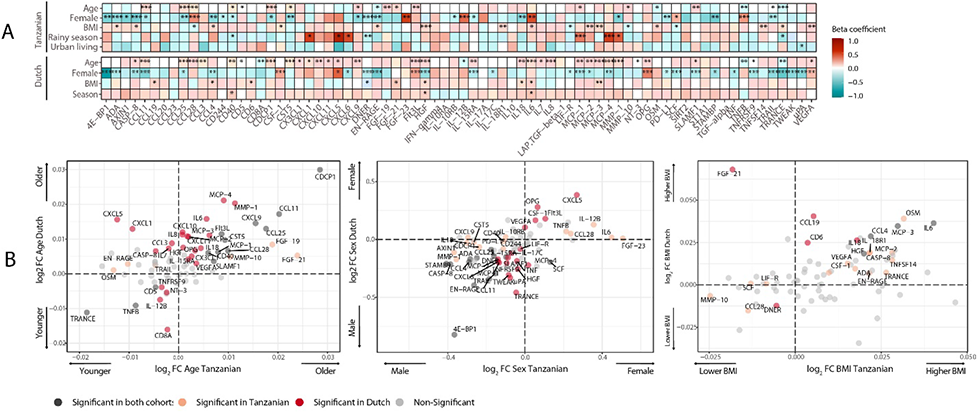
Associations of age, BMI, sex and seasonality with inflammatory proteins in the Dutch and Tanzanian participants. A) Heat maps showing the regression beta-coefficient of the plasma inflammatory proteins with host and environmental factors in the Dutch and Tanzanian cohorts. Dark red colors indicate higher concentrations of the inflammatory proteins with advanced age, higher BMI and higher in males, whereas turquoise color indicated lower concentration of inflammatory proteins with advanced age, higher BMI and higher concentrations in females in both cohorts. Depicted are p-values of the significant associations and results were declared significant after correcting for multiple testing using False discovered rate (FDR); p-value <0.05(*), <0.005(**), and <0.0001(***). B) Four-quadrant plot depicting the association between the inflammatory protein expression with either age, sex or BMI in the Dutch and Tanzania cohorts related to panel A

The most prominently downregulated proteins in Tanzanians were ST1A1 (sulfotransferase 1A1; log_2_ FC −0.8; *P*=2.2 x 10^-16^) and AXIN1 (axis inhibition protein 1; log_2_ FC −0.7; *P*=6.5 x 10^-18^). ST1A1 is a cytosolic sulfotransferase that catalyzes the sulfonation of endogenous and exogenous compounds (Wang, Cook, & Leyh, 2016). AXIN1 is a negative regulator of the Wnt/β-catenin signaling pathway (Kikuchi, 1999). This pathway is increasingly recognized to play an important role in inflammatory diseases, diabetes and cancer (Das, Das, Kalita, & Baro, 2021; Jridi, Canté-Barrett, Pike-Overzet, & Staal, 2021). Conversely, CDCP1 (CUB domain-containing protein 1), a transmembrane receptor that is a Wnt signaling promoter (He et al., 2020) was significantly up-regulated (log_2_ FC 0.7; *P*=1.8 x 10^-61^) in the Tanzanian cohort, suggesting enhanced activity of the Wnt/β-catenin signaling pathway in the Tanzanian participants. Finally, Tanzanians had lower levels of the CXC chemokine family members CXCL1, CXCL5, CXCL6, and CXCL8 (IL8). These chemokines mediate among others neutrophil trafficking (Palomino & Marti, 2015).

The Olink platform used in this study does not contain adipocytokines and provides relative, rather than absolute, protein concentrations. Therefore, we measured absolute concentrations of a selection of cytokines using an ELLA microfluidics platform, and concentrations of adipocytokines by ELISA. The data have been reported previously (Temba et al., 2022), and confirm that Tanzanian participants have significantly higher plasma concentrations of IL-1 receptor antagonist (IL-1Ra), IL-6, and IL-18, together with significantly higher leptin and lower adiponectin concentrations than Dutch participants after correcting for age, sex and BMI (Fig. 1c). In addition, Tanzanian females regardless of BMI had higher plasma concentrations of leptin compared to Dutch females (Supplemental Fig. 3) in line with the previous findings (Abbas, Lutale, & Ahren, 2004). There were no significant differences in plasma concentrations of resistin and alpha-1 antitrypsin (AAT).

### Associations between inflammation-related proteins and intrinsic and environmental factors

Next, we investigated associations between the inflammation-related proteins with host intrinsic factors such as age and sex, BMI and environmental exposures relevant to the Tanzanian setting. The analyses were variously corrected for age, sex and BMI to assess the impact of one specific factor. In high-income countries, age is a potent driver of immune variation with a shift toward a pro-inflammatory state (*25, 26*). In the Dutch participants, advancing age was indeed associated with an increase in inflammation-related proteins, including inflammatory cytokines (IL-6, IL-7, IL-18), IL-15RA, monocyte chemoattractant proteins (MCP2-4), matrix metalloprotease1 (MMP1), the chemokine IL-8 and hepatocyte growth factor (HGF) (Bielinski et al., 2017). In contrast, these significant associations were largely absent in the Tanzanians (Fig. 2 **&** Table S2). An exception was a strong significant positive association of advancing age with CDCP1, CCL11 and CCL25, which was also observed in the Dutch cohort. Overall, these results show that the association between advancing age and inflammatory markers is much weaker in the Tanzanians. Females in both cohorts overall had lower concentrations of inflammatory proteins than males (Fig. 2), which is consistent with our earlier findings that females had lower *ex vivo* cytokine responses (Temba et al., 2021; Ter Horst et al., 2016). Results also showed that Tanzanian females had significantly higher concentrations of TNF-beta (TNFB), IL12-beta and CCL28 than Tanzanian males.

Finally, since environmental exposures are potential drivers of inflammation, we determined the relationship between relevant exposures for Tanzanians and the proteome. Such exposures included type of toilet, exposure to wood smoke for cooking, farm-animal exposure, access to clean water, previous infections and prior use of antibiotics (Table 1). Our results did not show a significant association between inflammatory proteins with one of these exposures.

### Associations with food-derived metabolites and dietary habits

We recently reported that diet, and especially the transition between a rural traditional diet to an urban Western-type diet, had a major influence on *ex vivo* cytokine immune responses in the Tanzanian cohort (Temba et al., 2021). We postulated that diet also explained part of the variation in inflammatory proteins. To test this hypothesis, we selected food-derived metabolites (n=288) from an untargeted plasma metabolome, as previously described (Temba et al., 2021). Using these metabolites, we first performed unsupervised hierarchical clustering, which yielded two different clusters (food-metabolome clusters one and two) (Supplemental Fig. 4a). Weekly food consumption was associated with these food-metabolome clusters: participants in cluster one more frequently consumed ugali (a traditional porridge made from maize), plantain (cooking banana) and green vegetables, and less frequently rice and fried potato chips (Supplemental Fig. 4b). Next, we performed unsupervised clustering of the inflammatory proteome with age, sex, BMI, geolocation (i.e., rural vs. urban living), seasonality and the food-metabolome clusters as input variables. This analysis revealed two significant inflammatory proteome clusters: one with lower and one with higher expressed inflammatory proteins (Fig. 3). Participants belonging to food-metabolome cluster one (i.e., more ‘traditional’ Tanzanian diet) were overrepresented in the cluster with lower-expressed inflammatory proteins, whereas participants belonging to cluster two were overrepresented in the cluster of higher-expressed inflammatory proteins (Table S3). Other factors such as age, sex or season were not associated with the inflammatory clustering.

**Figure 3.**
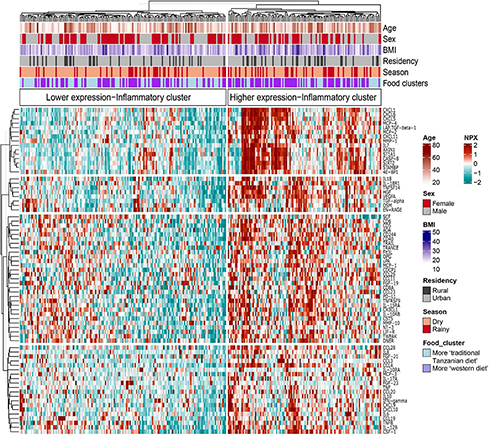
Association between food-derived plasma metabolites with inflammation-associated proteins. Unsupervised k-means clustering of individuals from the Tanzanian cohort (N= 318) according to the inflammatory proteins (N=72 inflammatory proteins). Data are shown as normalized protein expression (NPX). The color code indicates the relative expression of the inflammatory protein across the samples of the two compared groups. Dark red and turquoise colors indicate higher and lower expressions, respectively. Presented are annotations for age, sex, BMI, seasonality, geographical location (i.e rural vs. urban) and food-derived metabolite cluster. Results were considered significant after correcting for multiple testing using False discovered rate (FDR); p-value <0.05(*), <0.005(**), and <0.0001(***). Abbreviations: NXP; normalized protein expression; BMI; body mass index.

Next, we performed a correlation analysis to assess the relationship between diet-related metabolites and inflammatory proteins. Results show different negative associations between plant-derived polyphenols (apigenin, naringenin, cyanidin 3-(6-caffeoyl glucoside) 5-glucoside, licoagrodin, shoyuflavone C and phenolic acids such as gallic acid) and inflammatory proteins, particularly the MCP and CXCL families (Fig. 4). In contrast, positive associations were observed between inflammatory proteins, and especially members of the MCP and CXCL families, with plasma metabolites belonging to the following classes: carboxylic acids and derivatives (for example, aminobutanoic acid-ABA), organooxygen compounds (for example, triose), and prenol lipids such as resveratrol 4”-(6-) galloylglucoside (Fig. 4 **&** Table S4). The detailed classifications of various diet-derived metabolites and their correlations with various inflammatory proteins are presented in Table S4. Overall, these findings support the notion that a traditional plant-based Tanzanian diet in healthy Tanzanians has an important impact on circulating inflammatory proteins.

**Figure 4.**
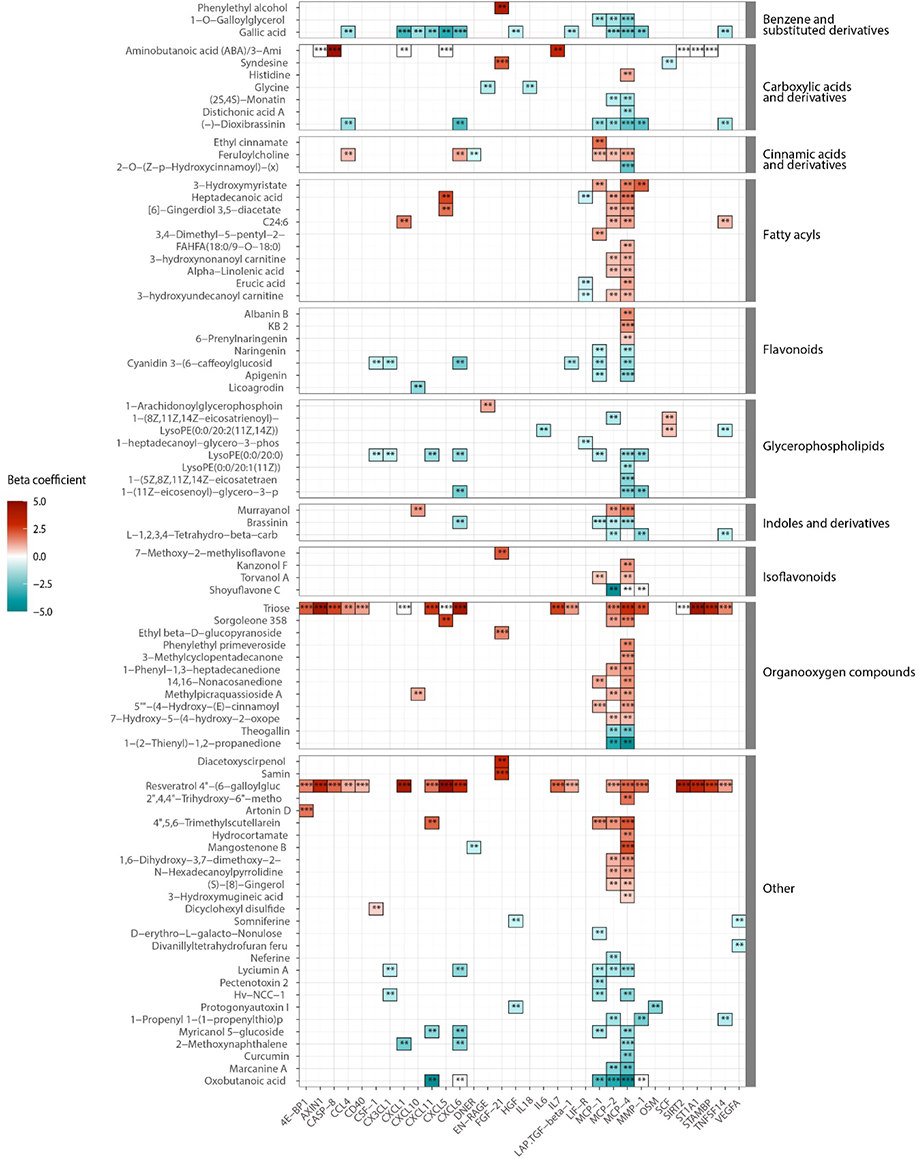
Association between inflammatory proteins and the food-derived metabolites within the Tanzanian cohort. Heat map showing associations between inflammatory proteins and food-derived metabolites within the Tanzanian participants. Depicted are β coefficients of the multiple linear regression model, including age and sex as covariates. Red and turquoise colors indicate positive and negative correlations, respectively. results were declared significant after correcting for multiple testing using False discovered rate (FDR); p-value <0.05(*), <0.005(**), and <0.0001(***).

## DISCUSSION

Genetic and environmental variations, including diet, lifestyle and infectious diseases burden, are important for modulating immune responses and may result in differences in immune phenotypes across populations. Our analysis of the inflammatory proteome shows that healthy Tanzanians have a more prominent pro-inflammatory phenotype compared to healthy Dutch individuals. Among the Tanzanians, food-derived metabolites were identified as an important driver of variation in inflammation-related proteins, emphasizing the potential importance of lifestyle changes.

In the same European and African cohorts, we recently showed remarkable differences in the genetic regulation of immune responses, with ancestry-specific pathways regulating induced cytokine responses and significant enrichment of the interferon pathway in the Tanzanians (Boahen et al., 2022). From a historical perspective, the natural selection of genotypes that mediate a strong inflammatory response offers an obvious advantage in areas with a high infectious diseases burden such as sub-Saharan Africa. In addition, epigenetic regulation as a conduit that mirrors environmental exposure (e.g., diet, lifestyle or exposure to infectious diseases) may also promote an inflammatory signature. However, while a strong inflammatory response can be advantageous in the host’s response against infections, it may become disadvantageous when the environment changes with a reduction in the infectious burden and a shift toward an unhealthy lifestyle. Many areas in sub-Saharan Africa are currently witnessing such changes and this is one of the important drivers of the rapid increase in non-communicable diseases and other inflammation-associated pathologies.

A healthy diet is an important component of a healthy lifestyle. In the present study, we show that Tanzanians who consumed more traditional staple foods, such as ugali (porridge made from maize or millet) or plantain, as well as green vegetables, had lower concentrations of inflammation-related proteins. We also observed several significant associations between food-derived metabolites and inflammatory proteins. Specifically, plant-derived polyphenols were negatively associated with members of the MCP and CXCL families. Especially gallic acid, a natural phenic acid with different health benefits, which is present in mangos and other edible plants (Kim et al., 2021), had different negative associations with immune-derived proteins. These data confirm our earlier findings that diet has an important impact on cytokine responses (Temba, 2021 #287) as well as plasmatic coagulation (Temba et al., 2022). Traditional foods, particularly plant-based diets, are increasingly recognized for their cardiovascular health benefits (Hemler & Hu, 2019), and our findings support the notion that promoting traditional diets may be a viable public health strategy for curbing the NCDs epidemic. Other environmental exposures may potentially confound the diet-inflammation associations, for example, individuals with a traditional diet may also use more frequently a pit latrine or smoky fuels and may have more infections, but we found no significant associations between inflammatory proteins and any of these other environmental exposures.

An interesting observation was the limited number of associations between advancing age and inflammation-related proteins in the Tanzanians. In historically wealthy countries, aging is associated with a blunted response of immune cells to stimulation, while having a chronic low-inflammation state (Guarner & Rubio-Ruiz, 2015; Liberale, Montecucco, Tardif, Libby, & Camici, 2020), as also observed in Dutch in this study. Cytokine responses decline in the Tanzanians with advancing age (Temba et al., 2021), similar to Europeans, but the reasons why systemic inflammation does not increase remain uncertain. Possible explanations are the already present inflammatory signature at a younger age, possibly because of higher cumulative exposure to infections or other environmental insults, or those older participants in Tanzania manage to control the inflammatory state by a healthier lifestyle.

The two most upregulated proteins in the Tanzanians, 4E-BP1 and FGF21, are regulators of metabolism. FGF21 has received much attention recently as an endocrine regulator of glucose, lipid and energy metabolism (Hill et al., 2018). FGF21 is an insulin sensitizer, and its secretion is increased by a high intake of carbohydrates and low dietary protein intake (Lundsgaard et al., 2017; Maekawa et al., 2017). FGF21 inhibits age-associated metabolic syndrome and protects against diabetic cardiomyopathy (Yan et al., 2021). FGF21, therefore, acts differently from classic energy balance signals like leptin (Hill et al., 2018). Traditional Tanzanian diets are high in carbohydrates and low in proteins, which may explain the high FGF21 among the Tanzanians. FGF21 was also reported to be higher in obesity (X. Zhang et al., 2008), but we did not find an association with BMI. Tanzanians also had higher leptin and lower adiponectin concentrations than the Dutch. This is in line with the results of an earlier study among non-obese and obese subjects with type 2 diabetes in Tanzania and Sweden, which showed that leptin concentrations were 50% higher in the Tanzanians (Abbas et al., 2004). This supports the importance of potential ethnic differences in metabolic profiles and adipocytokines (Mente et al., 2010). To our knowledge, this study is the first to compare FGF21 levels between healthy individuals from sub-Saharan Africa and Europe.

Another interesting finding was the significant differences in CDCP1 and AXIN1, pointing towards an enhanced activity of the Wnt/β-catenin pathway in the Tanzanians. The Wnt/β-catenin pathway is a key regulator of inflammation, playing a role in both the inflammatory and anti-inflammatory pathways (Ma & Hottiger, 2016). Dysregulated activation of the Wnt/β-catenin pathway is increasingly recognized to play a role in the pathogenesis of chronic inflammatory diseases, metabolic inflammatory diseases and cancer (Jridi et al., 2021). Mice fed on a high-fat Western-type diet expressed high concentrations of Wnt2 protein in atherosclerotic lesions, suggesting that the Wnt/β-catenin pathway also contributes to atherosclerosis (J. Zhang et al., 2021).

To summarize, our findings reveal significant differences in inflammatory and metabolic proteins and pathways between healthy individuals living in East Africa and individuals living in Western Europe. This is especially important in light of the current epidemiological transition and lifestyle changes in sub-Saharan Africa, which coincides with a sharp increase in non-communicable diseases in the region. Our study also endorses the importance of including underrepresented populations in systems-immunology studies.

## Material and Methods

### Study Design and Population

The present study used samples from two cross-sectional cohorts of healthy volunteers: the 300-Tanzania-FG (300TZFG) and the Dutch 500FG. Both cohorts were enrolled within the Human Functional Genomics Project (https://www.humanfunctionalgenomics.org). The demographic characteristics of both cohorts have been described previously (Temba et al., 2021; Ter Horst et al., 2016). Briefly, the 300TZFG cohort consists of 323 healthy Tanzanian individuals aged between 18 and 65 years residing in the Kilimanjaro region in Northern Tanzania. The cohort was enrolled between March and December 2017. Exclusion criteria were participants with any acute or chronic disease, use of antibiotics or anti-malaria medication in the three months before blood sampling, tuberculosis in the past year, a blood pressure ≤ 90/60mmHg or ≥140/90mmHg, or random blood glucose >8.0 mmol/L. Pregnant, postpartum, or breastfeeding females were excluded. The 500FG cohort consists of 534 Dutch individuals of Western-European background, aged 18 years and older. Data was collected between August 2013 and December 2014 at the Radboud university medical center (Radboudumc) in the Netherlands. Exclusion criteria were: the use of any medication in the past month and acute or chronic diseases at the time of blood sampling. Pregnant, postpartum, or breastfeeding females were excluded.

### Sample collection and preparation

The current study is part of the Human Functional Genomics Project (HFGP; humanfunctionalgenomics.org), which employs standardized procedures for sample collection, handling, and pre-processing. Blood was obtained in the morning via antecubital puncture into ethylenediaminetetraacetic acid (EDTA) tubes (Monoject^TM^; Covidien, Ireland). Within two to three hours after blood collection, plasma was collected by centrifugation at 3800rpm for 8 minutes at room temperature. The obtained plasma were stored at −80°C, as recommended by the ISBER biobanking organization (Garcia, Bracci, Guevarra, & Sieffert, 2014). Plasma samples for the Tanzania cohort were shipped to the Netherlands on dry ice.

### Inflammatory proteome

Plasma proteins were measured with the Olink 92 Inflammation panel using proximity extension technology (Olink® Proteomics AB, Uppsala, Sweden)(Assarsson et al., 2014). This panel includes 92 inflammation-related proteins. This assay utilizes the binding of target proteins by paired oligonucleotide antibody probes, followed by hybridization and amplification. Data are reported as normalized protein expression values (NPX), which is an arbitrary unit in a Log2 scale that is calculated from normalized Ct values. Validation data of the assay are available on the Olink website (www.olink.com). All samples were measured in the same batch in a single run. Proteins were excluded from analysis when values were both below the detection limit in more than 25% of all samples. Plasma samples from the Tanzanian and Dutch cohorts were on their first and second freeze-thawed cycles, respectively. Pre-analytical processing such as freeze-thawed cycles and storage time has limited influence on the measured proteins reported in this study (Enroth, Hallmans, Grankvist, & Gyllensten, 2016; Lee, Kim, & Shin, 2015; Shen et al., 2018).

### Measurement of the circulating inflammatory mediators

Plasma concentrations of the cytokines IL-6, IL-1β, IL-1 receptor antagonist (IL-1Ra) and IL-18 (lot number Bio-Tech/R&D; SPCKC-PS-001559) and IL-18 binding protein (IL-18BPa) (lot number Bio-Tech/R&D; SPCKB-PS-000502) were measured in EDTA plasma using the Simple Plex^TM^ cartridges run on the Ella^TM^ platform (Protein Simple, San Jose, USA) following the manufacturer’s instructions.

### Plasma metabolome

Plasma samples of the Tanzanian cohort were measured using the untargeted metabolomics workflow by General Metabolics (Boston, MA) with procedures as previously described (Fuhrer, Heer, Begemann, & Zamboni, 2011). In short, metabolites were measured by a high throughput mass spectrometry technique using the Agilent Series 1100 LC pump coupled to a Gerstel MPS2 autosampler and the Agilent 6520 Series Quadrupole Time-of-flight mass spectrometer (Agilent, Santa Clara, CA). The selection of food-derived metabolites was performed based on the ontology given in the HMDB (https://www.hmdb.ca/) as described previously (Temba et al., 2021).

### Ethical statement

The 300TZFG study was approved by the Ethical Committees of the Kilimanjaro Christian Medical University College (CRERC) (No 2443) and the National Institute for Medical Research (NIMR/HQ/R.8a/Vol. IX/2290 and NIMR/HQ/R.8a/Vol.IX/3318) in Tanzania. The 500FG cohort study was approved by the Ethical Committee of the Radboud University Medical Centre Nijmegen, the Netherlands (NL42561.091.12, 2012/550). Subject recruitment and experimental procedures were conducted according to the principles mentioned in the Declaration of Helsinki. Written informed consent was obtained from all subjects.

### Statistical analysis

The proteomic data from the Dutch and Tanzanian cohorts were normalized using inter-plate controls for batch variation correction and presented in the log_2_ scale. Data values below the limit of detection (LOD) were handled using the actual measured values to increase the statistical power and give a complete data distribution. Outliers detection were done using principal component analysis (PCA) in which data points that fall in more than 3 standard deviations from the mean of principal component one (PC1) and two (PC2) were excluded. This pre-analytical process led to exclusion of 12 participants (N=7 Dutch and N=5 Tanzanians) as potential outliers. Proteins with >25% data values below LOD in both cohorts were excluded (N=18 in the Duch and N=20 in the Tanzania cohorts), leaving 74 and 72 inflammatory proteins for the downstream data analysis in the Dutch and Tanzanian cohorts respectively. In total, 416 Dutch and 318 Tanzania participants were available after the pre-analytical process. Details of preanalytical steps for both cohorts are described in Supplementary figure 1.

To analyze the similarities and dissimilarities between the samples, unsupervised PCA was performed using ‘prcomp’ function in R package. Heatmap of unsupervised hierarchical clustering (k-nearest neighbors with 100 repetitions) of the samples was generated using ‘ComplexHeatmap’ R package by calculating the matrix of Euclidean distances from the log_2_ NPX value. Limma (linear models for microarray data) R package was used for differential expression analysis of plasma inflammatory proteins between cohorts. Proteins with adjusted P-value (FDR) < 0.05 were selected as significantly differentially expressed (DE). Proteins with a positive and negative value of log_2_-fold-change were considered as upregulated and downregulated, respectively.

### Data availability

Anonymized metadata of the Tanzanian participants and the circulating inflammation markers are available in an open access registry (DANS registry; https://doi.org/10.17026/dans-xgx-zuht) Untargeted plasma metabolome data have been deposited to the EMBL-EBI MetaboLights database (http://www.ebi.ac.uk/metabolights/); study identifier MTBLS2267. The source data of the proteomics analysis are provided in Supplemental Table 5. Publicly available databases used for this study include KEGG (https://www.genome.jp/kegg/), HMDB (https://www.hmdb.ca/) and ChEBI (https://ebi.ac.uk/chebi/). All other data is available in the main text and supplementary materials.

## Acknowledgments

The authors wish to express their gratitude to all volunteers in the Human Functional Genomics in Tanzania and Dutch cohorts for their participation. We would like to thank **J. Njau, J. Kwayu**, and **E. Kimaro** for assistance with sample collection, **H. Lemmers**, and **H. Toenhake-Dijkstra** for assistance with laboratory analysis, and **M. Miclaus** for assistance with pre-processing metabolome data.

## Author contributions

Q.d.M., A.V., M.G.N., L.A.B.J., R.K., J.L.S., and B.T.M contributed to the conceptualization, study design, data interpretation and lead the project; G.S.T., V.K., R.K., and B.T.M., contributed to participants recruitment, data collection and laboratory analyses; G.S.T and N.V performed statistical analysis and graphics of the proteome, inflammatory and metabolome data; G.S.T and Q.d.M. wrote the original draft of the manuscript; and G.S.T., N.V., T.P., V.K., B.T.M., R.K., L.A.B.J., J.L.S., A.V., M.G.N., and Q.d.M., contributed in writing-review and editing the manuscript.

## Supplemental Figures and Tables

**Supplemental Fig. 1:**
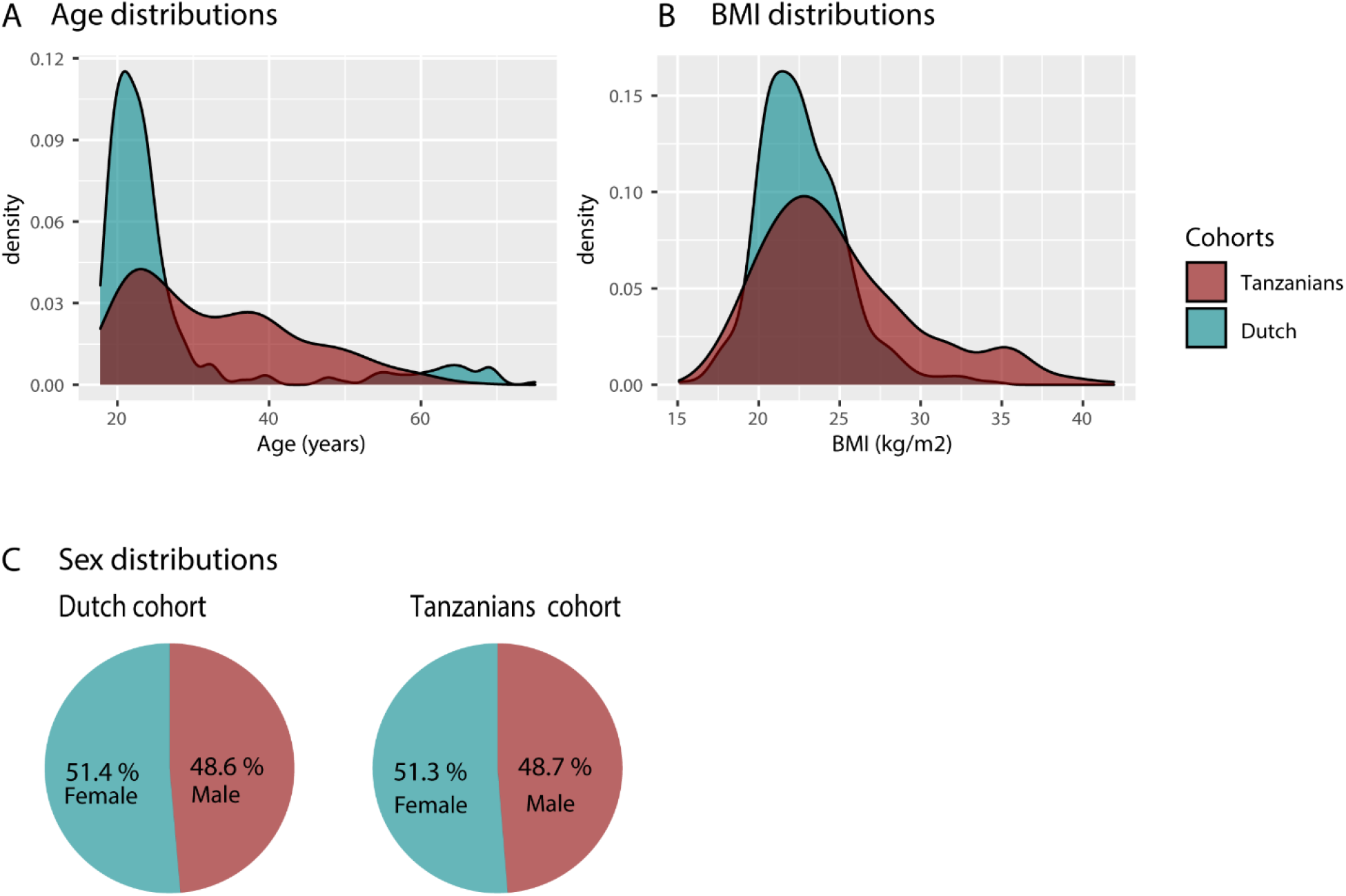
Histograms and pie charts showing the distribution of cohorts characteristics comparing Tanzanian and Dutch samples. Histograms **A)** and **B)** show the age and BMI distributions of Tanzanians and Dutch subjects, respectively) A pie chart comparing the gender distributions of Tanzanians and the Dutch cohort.

**Supplemental Fig. 2:**
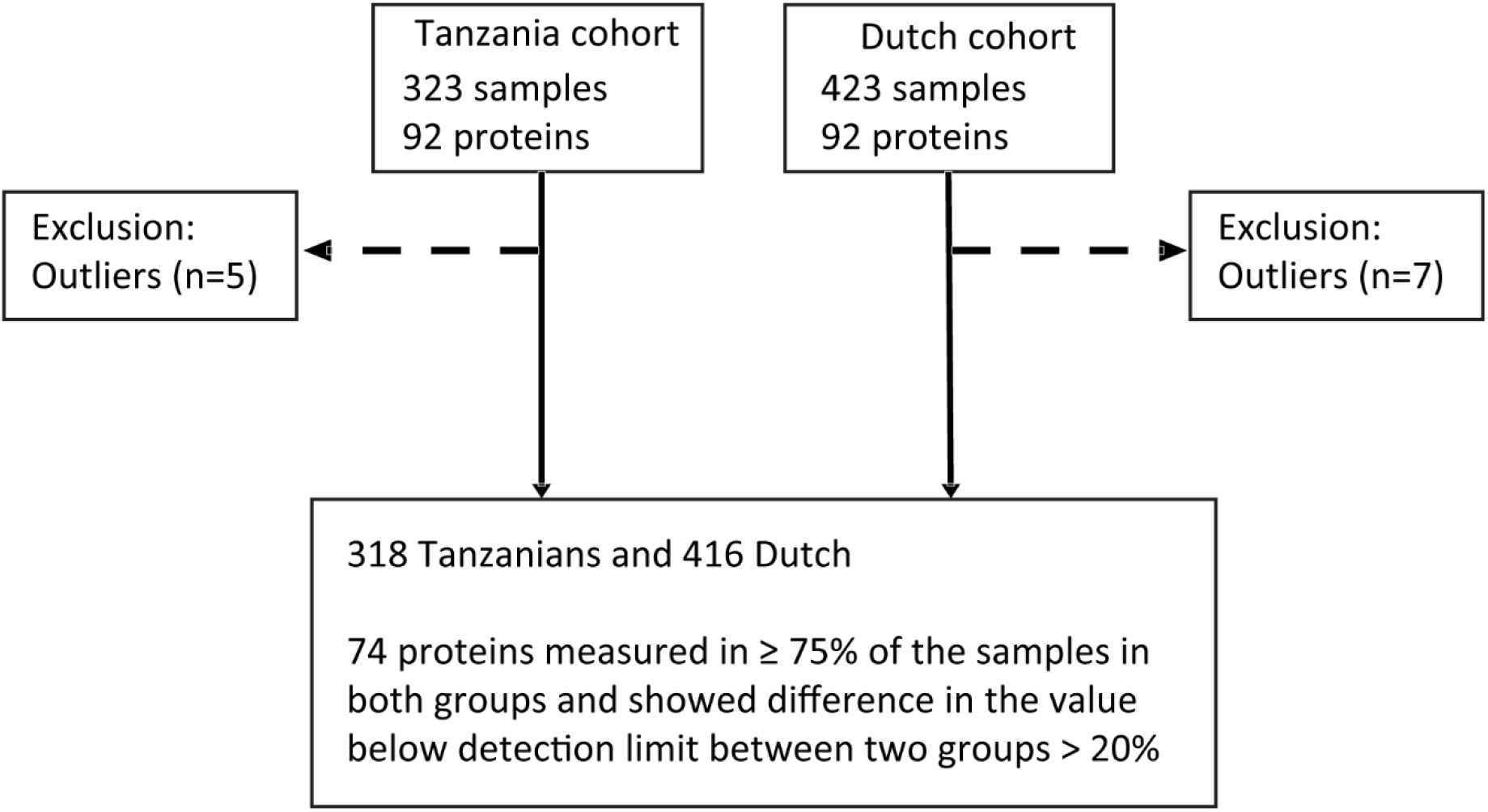
Schematic diagram showing the sample preprocessing according to the measured inflammatory protein in both cohorts

**Supplemental Fig. 3:**
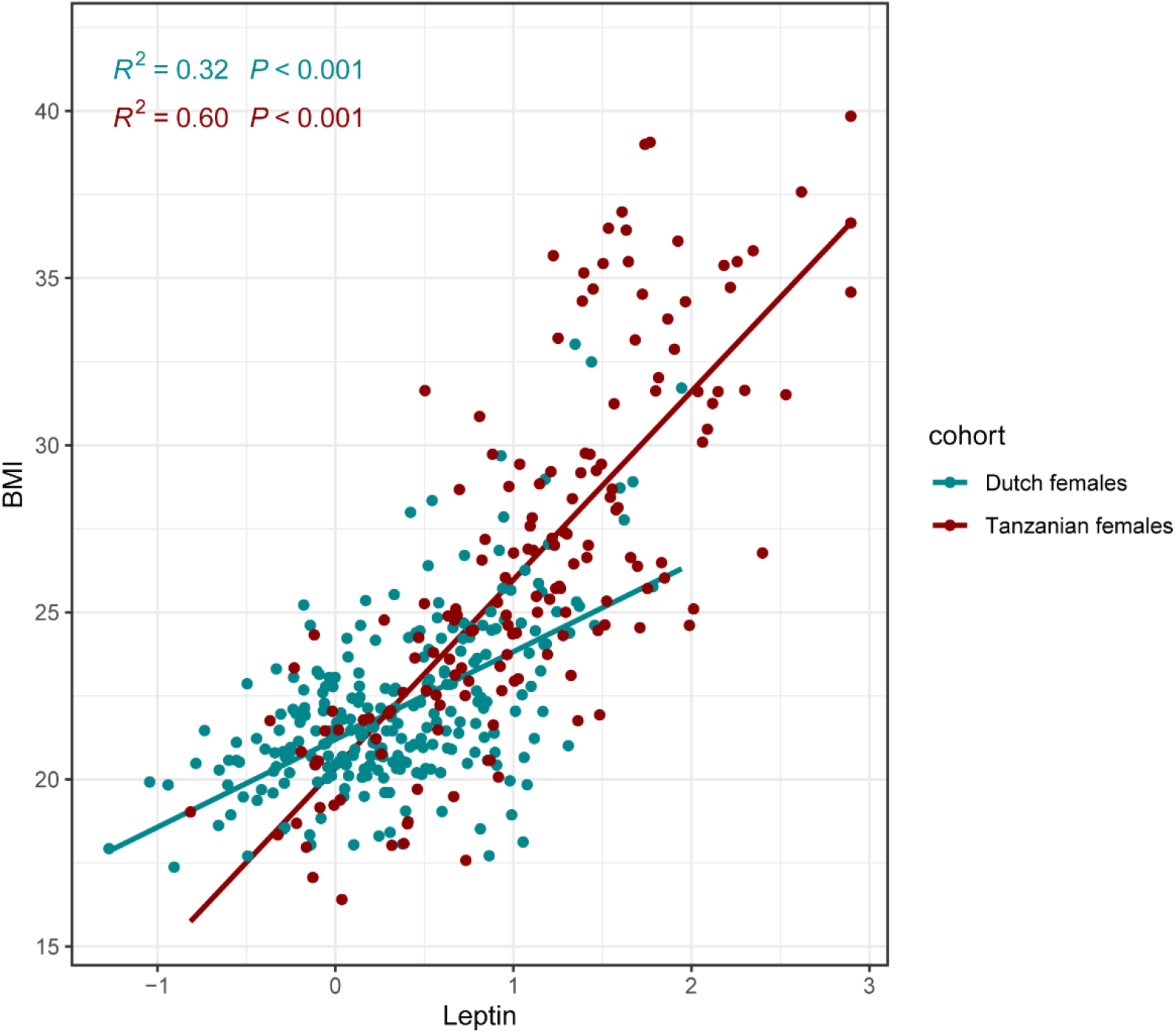
Scatter plot showing the association of plasma concentration of leptin with BMI in Tanzanian females compared to Dutch females. The comparison was performed using linear regression (unadjusted) on the inversed ranked based values of leptin concentration.

**Supplemental Fig. 4:**
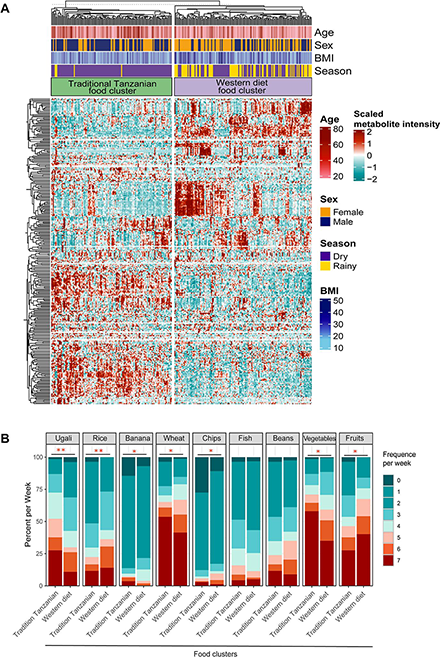
Food-metabolome clusters and their associations with weekly food consumption within the Tanzanian cohort. **A)** Heat map displaying unsupervised k-means clustering of individuals from the Tanzanian cohort (N = 318) based on food-derived metabolites (N = 298 food-derived metabolites). Food-derived metabolites from the untargeted plasma metabolome were clustered, which yielded two clusters (traditional Tanzanian food cluster and Western diet food cluster). Red and blue colors indicate higher and lower metabolite intensity, respectively. Presented are annotations for age, sex, BMI, and seasonality. **B)** Weekly food consumption of various foods across both metabolic clusters. Differences in food consumption frequency categories were tested using the chi-squared test. The colors red and blue represent higher and lower weekly consumption frequencies of specific foods, respectively; p-values 0.05 (*), 0.005 (**), and 0.0001 (***).BMI stands for body mass index.

**Table S1.** Differential expression analysis 300TZ vs 500FG cohort

| Proteins | av_expr | Log2FC (Tanzanian/Dutch) | pvalue | fdr |
| --- | --- | --- | --- | --- |
| 4E-BP1 | 6,596 | 1,875 | 1,06587E-61 | 1,31458E-60 |
| ADA | 5,153 | 0,528 | 2,7438E-52 | 2,90059E-51 |
| AXIN1 | 3,036 | -0,728 | 2,19745E-18 | 6,50445E-18 |
| CASP-8 | 3,036 | -0,432 | 1,09865E-19 | 3,53478E-19 |
| CCL11 | 5,900 | 0,679 | 2,95603E-64 | 7,29154E-63 |
| CCL19 | 8,580 | -0,039 | 0,47982057 | 0,514590176 |
| CCL20 | 6,296 | 0,477 | 8,82426E-13 | 1,92058E-12 |
| CCL23 | 10,041 | -0,328 | 3,98093E-18 | 1,09107E-17 |
| CCL25 | 5,662 | 0,309 | 1,83527E-13 | 4,11546E-13 |
| CCL28 | 1,718 | 0,396 | 2,19671E-39 | 1,62556E-38 |
| CCL3 | 3,380 | 0,557 | 2,92987E-10 | 5,55924E-10 |
| CCL4 | 4,986 | 0,358 | 1,34199E-15 | 3,4244E-15 |
| CD244 | 7,036 | 0,021 | 0,45012628 | 0,489843304 |
| CD40 | 10,998 | -0,212 | 2,66298E-12 | 5,63031E-12 |
| CD5 | 5,084 | 0,282 | 1,45729E-31 | 8,29532E-31 |
| CD6 | 5,026 | 0,351 | 1,22626E-21 | 4,53716E-21 |
| CD8A | 10,299 | 0,568 | 2,10053E-15 | 5,18131E-15 |
| CDCP1 | 1,268 | 0,732 | 9,84134E-63 | 1,82065E-61 |
| CSF-1 | 9,623 | 0,024 | 0,177665233 | 0,212052052 |
| CST5 | 5,228 | 0,484 | 6,25405E-34 | 3,85666E-33 |
| CX3CL1 | 5,801 | -0,015 | 0,622772179 | 0,640071406 |
| CXCL1 | 8,300 | -0,412 | 2,29827E-05 | 3,61855E-05 |
| CXCL10 | 8,337 | -0,236 | 4,33405E-05 | 6,68165E-05 |
| CXCL11 | 7,016 | -0,042 | 0,557401936 | 0,580954131 |
| CXCL5 | 9,100 | -0,548 | 1,0954E-05 | 1,76217E-05 |
| CXCL6 | 8,019 | -0,554 | 2,24875E-14 | 5,20022E-14 |
| CXCL9 | 6,578 | 0,111 | 0,076518212 | 0,097626684 |
| DNER | 9,268 | 0,398 | 1,34737E-82 | 9,97056E-81 |
| EN-RAGE | 1,757 | 0,380 | 1,86908E-14 | 4,46166E-14 |
| FGF-19 | 8,168 | 0,195 | 0,005867961 | 0,008350561 |
| FGF-21 | 4,470 | 1,325 | 2,38017E-31 | 1,25809E-30 |
| FGF-23 | 2,035 | 0,052 | 0,366680053 | 0,408915624 |
| Flt3L | 8,594 | -0,120 | 7,99968E-05 | 0,000120811 |
| HGF | 7,444 | 0,246 | 2,92717E-18 | 8,33117E-18 |
| IFN-gamma | 5,931 | -0,053 | 0,531423753 | 0,561790825 |
| IL-10RA | 1,202 | 0,064 | 0,286528227 | 0,326201366 |
| IL-10RB | 5,717 | 0,033 | 0,137157208 | 0,166387433 |
| IL-12B | 4,981 | -0,074 | 0,092338531 | 0,113884188 |
| IL-12B | 4,981 | -0,074 | 0,092338531 | 0,113884188 |
| IL-15RA | 1,106 | -0,107 | 3,12389E-08 | 5,63824E-08 |
| IL-17A | 1,749 | 0,709 | 1,43118E-34 | 9,62792E-34 |
| IL-17C | 1,177 | 0,713 | 3,0894E-76 | 1,14308E-74 |
| IL-18R1 | 7,612 | -0,011 | 0,722011984 | 0,731902559 |
| IL10 | 3,493 | 0,397 | 4,32004E-12 | 8,88009E-12 |
| IL18 | 7,416 | 0,396 | 1,84574E-18 | 5,69104E-18 |
| IL6 | 1,390 | 0,193 | 0,001294948 | 0,001916523 |
| IL7 | 2,779 | -0,368 | 7,9284E-11 | 1,54395E-10 |
| IL8 | 4,203 | -0,289 | 1,48778E-06 | 2,50218E-06 |
| LAP.TGF-beta-1 | 7,126 | -0,199 | 1,82213E-07 | 3,21042E-07 |
| LIF-R | 3,759 | 0,357 | 3,62208E-62 | 5,36068E-61 |
| MCP-1 | 10,375 | -0,445 | 1,36861E-30 | 6,75183E-30 |
| MCP-2 | 7,256 | 0,499 | 2,15356E-23 | 8,38757E-23 |
| MCP-3 | 0,573 | 0,609 | 8,98684E-43 | 7,38918E-42 |
| MCP-4 | 11,402 | -0,100 | 0,064788541 | 0,084111439 |
| MMP-1 | 8,996 | -0,372 | 1,89878E-06 | 3,12243E-06 |
| MMP-10 | 8,567 | 0,235 | 1,26357E-06 | 2,17451E-06 |
| NT-3 | 3,061 | -0,098 | 0,009930266 | 0,0138649 |
| OSM | 2,845 | 0,613 | 3,46374E-21 | 1,22056E-20 |
| PD-L1 | 6,595 | -0,078 | 0,003293161 | 0,004778312 |
| SCF | 9,155 | 0,185 | 5,6904E-10 | 1,05272E-09 |
| SIRT2 | 4,459 | -0,198 | 0,031175292 | 0,041944939 |
| SLAMF1 | 1,389 | 0,278 | 2,015E-27 | 9,31936E-27 |
| ST1A1 | 4,094 | -0,772 | 8,21151E-17 | 2,17019E-16 |
| STAMBP | 5,359 | -0,143 | 0,023265062 | 0,031881751 |
| TGF-alpha | 2,925 | 0,383 | 5,93976E-52 | 5,49428E-51 |
| TNF | 2,646 | 0,479 | 8,21195E-20 | 2,7622E-19 |
| TNFB | 4,819 | 0,044 | 0,201404932 | 0,236570872 |
| TNFRSF9 | 6,993 | 0,036 | 0,223880475 | 0,258861799 |
| TNFSF14 | 3,132 | 0,063 | 0,087857702 | 0,110194406 |
| TRAIL | 7,429 | 0,266 | 9,26858E-26 | 4,03456E-25 |
| TRANCE | 4,783 | -0,046 | 0,370234416 | 0,408915624 |
| TWEAK | 9,311 | 0,294 | 2,99206E-25 | 1,23007E-24 |
| uPA | 9,893 | 0,218 | 5,83238E-11 | 1,16648E-10 |
| VEGFA | 9,554 | -0,007 | 0,759842371 | 0,759842371 |

**Table S2.** Multivariate linear regression analysis for factors affecting protein expression in Tanzanian cohort

| Proteins | Covariate | Estimate | Std. Error | t value | pvalue | fdr |
| --- | --- | --- | --- | --- | --- | --- |
| 4E-BP1 | Age | -0,0110 | 0,0045 | -2,4367 | 0,0154 | 0,0670 |
| 4E-BP1 | Female | -0,3651 | 0,0919 | -3,9715 | 0,0001 | 0,0012 |
| 4E-BP1 | BMI | 0,0185 | 0,0101 | 1,8282 | 0,0685 | 0,1964 |
| 4E-BP1 | Residency_AreaUrban | 0,0551 | 0,1120 | 0,4921 | 0,6230 | 0,7841 |
| 4E-BP1 | Seasonality_Rainy | 0,1432 | 0,0950 | 1,5080 | 0,1326 | 0,3170 |
| ADA | Age | -0,0055 | 0,0028 | -1,9776 | 0,0489 | 0,1588 |
| ADA | Female | -0,2835 | 0,0570 | -4,9718 | 0,0000 | 0,0000 |
| ADA | BMI | 0,0174 | 0,0063 | 2,7777 | 0,0058 | 0,0330 |
| ADA | Residency_AreaUrban | -0,1337 | 0,0695 | -1,9248 | 0,0552 | 0,1709 |
| ADA | Seasonality_Rainy | -0,0336 | 0,0589 | -0,5711 | 0,5683 | 0,7605 |
| AXIN1 | Age | -0,0083 | 0,0062 | -1,3467 | 0,1791 | 0,3743 |
| AXIN1 | Female | -0,3679 | 0,1264 | -2,9113 | 0,0039 | 0,0267 |
| AXIN1 | BMI | 0,0305 | 0,0139 | 2,1910 | 0,0292 | 0,1080 |
| AXIN1 | Residency_AreaUrban | -0,2318 | 0,1539 | -1,5062 | 0,1330 | 0,3170 |
| AXIN1 | Seasonality_Rainy | -0,0027 | 0,1305 | -0,0204 | 0,9837 | 0,9918 |
| CASP-8 | Age | -0,0046 | 0,0036 | -1,2783 | 0,2021 | 0,4086 |
| CASP-8 | Female | -0,3861 | 0,0739 | -5,2241 | 0,0000 | 0,0000 |
| CASP-8 | BMI | 0,0215 | 0,0081 | 2,6435 | 0,0086 | 0,0449 |
| CASP-8 | Residency_AreaUrban | -0,0502 | 0,0900 | -0,5573 | 0,5777 | 0,7641 |
| CASP-8 | Seasonality_Rainy | -0,0569 | 0,0764 | -0,7457 | 0,4564 | 0,6701 |
| CCL11 | Age | 0,0203 | 0,0026 | 7,7792 | 0,0000 | 0,0000 |
| CCL11 | Female | -0,1850 | 0,0532 | -3,4780 | 0,0006 | 0,0058 |
| CCL11 | BMI | -0,0111 | 0,0059 | -1,8880 | 0,0600 | 0,1808 |
| CCL11 | Residency_AreaUrban | -0,0119 | 0,0648 | -0,1830 | 0,8549 | 0,9390 |
| CCL11 | Seasonality_Rainy | 0,1023 | 0,0550 | 1,8616 | 0,0636 | 0,1868 |
| CCL19 | Age | -0,0025 | 0,0043 | -0,5715 | 0,5681 | 0,7605 |
| CCL19 | Female | -0,0078 | 0,0877 | -0,0885 | 0,9295 | 0,9688 |
| CCL19 | BMI | 0,0053 | 0,0097 | 0,5525 | 0,5810 | 0,7641 |
| CCL19 | Residency_AreaUrban | 0,0965 | 0,1068 | 0,9031 | 0,3672 | 0,5830 |
| CCL19 | Seasonality_Rainy | -0,0631 | 0,0906 | -0,6969 | 0,4864 | 0,7003 |
| CCL20 | Age | -0,0076 | 0,0051 | -1,5084 | 0,1325 | 0,3170 |
| CCL20 | Female | -0,0159 | 0,1034 | -0,1542 | 0,8775 | 0,9487 |
| CCL20 | BMI | -0,0021 | 0,0114 | -0,1887 | 0,8504 | 0,9390 |
| CCL20 | Residency_AreaUrban | -0,3156 | 0,1259 | -2,5068 | 0,0127 | 0,0584 |
| CCL20 | Seasonality_Rainy | 0,1651 | 0,1068 | 1,5463 | 0,1230 | 0,2995 |
| CCL23 | Age | 0,0035 | 0,0029 | 1,2000 | 0,2311 | 0,4453 |
| CCL23 | Female | -0,1330 | 0,0594 | -2,2398 | 0,0258 | 0,0984 |
| CCL23 | BMI | 0,0047 | 0,0065 | 0,7207 | 0,4717 | 0,6844 |
| CCL23 | Residency_AreaUrban | -0,0826 | 0,0723 | -1,1420 | 0,2543 | 0,4674 |
| CCL23 | Seasonality_Rainy | -0,0856 | 0,0613 | -1,3948 | 0,1641 | 0,3571 |
| CCL25 | Age | 0,0176 | 0,0029 | 6,0714 | 0,0000 | 0,0000 |
| CCL25 | Female | -0,2490 | 0,0593 | -4,2006 | 0,0000 | 0,0006 |
| CCL25 | BMI | -0,0113 | 0,0065 | -1,7294 | 0,0847 | 0,2239 |
| CCL25 | Residency_AreaUrban | -0,0393 | 0,0722 | -0,5438 | 0,5870 | 0,7641 |
| CCL25 | Seasonality_Rainy | -0,0320 | 0,0612 | -0,5227 | 0,6015 | 0,7728 |
| CCL28 | Age | 0,0099 | 0,0023 | 4,3804 | 0,0000 | 0,0003 |
| CCL28 | Female | 0,2375 | 0,0463 | 5,1300 | 0,0000 | 0,0000 |
| CCL28 | BMI | -0,0137 | 0,0051 | -2,6886 | 0,0076 | 0,0411 |
| CCL28 | Residency_AreaUrban | 0,0027 | 0,0564 | 0,0474 | 0,9622 | 0,9835 |
| CCL28 | Seasonality_Rainy | 0,0225 | 0,0478 | 0,4704 | 0,6384 | 0,7900 |
| CCL3 | Age | -0,0014 | 0,0093 | -0,1473 | 0,8830 | 0,9487 |
| CCL3 | Female | 0,0722 | 0,1899 | 0,3801 | 0,7041 | 0,8377 |
| CCL3 | BMI | 0,0290 | 0,0209 | 1,3885 | 0,1660 | 0,3575 |
| CCL3 | Residency_AreaUrban | -0,3653 | 0,2312 | -1,5797 | 0,1152 | 0,2900 |
| CCL3 | Seasonality_Rainy | -0,0086 | 0,1961 | -0,0441 | 0,9649 | 0,9835 |
| CCL4 | Age | -0,0016 | 0,0034 | -0,4818 | 0,6303 | 0,7852 |
| CCL4 | Female | -0,2893 | 0,0695 | -4,1659 | 0,0000 | 0,0007 |
| CCL4 | BMI | 0,0220 | 0,0076 | 2,8746 | 0,0043 | 0,0282 |
| CCL4 | Residency_AreaUrban | -0,0643 | 0,0846 | -0,7600 | 0,4479 | 0,6602 |
| CCL4 | Seasonality_Rainy | 0,1761 | 0,0717 | 2,4538 | 0,0147 | 0,0647 |
| CD244 | Age | 0,0000 | 0,0021 | -0,0016 | 0,9987 | 0,9987 |
| CD244 | Female | -0,1125 | 0,0430 | -2,6167 | 0,0093 | 0,0472 |
| CD244 | BMI | 0,0027 | 0,0047 | 0,5605 | 0,5756 | 0,7641 |
| CD244 | Residency_AreaUrban | -0,0125 | 0,0524 | -0,2379 | 0,8121 | 0,9189 |
| CD244 | Seasonality_Rainy | -0,0607 | 0,0444 | -1,3676 | 0,1724 | 0,3625 |
| CD40 | Age | 0,0059 | 0,0019 | 3,0564 | 0,0024 | 0,0196 |
| CD40 | Female | -0,1119 | 0,0398 | -2,8146 | 0,0052 | 0,0310 |
| CD40 | BMI | 0,0104 | 0,0044 | 2,3809 | 0,0179 | 0,0727 |
| CD40 | Residency_AreaUrban | -0,0510 | 0,0484 | -1,0538 | 0,2928 | 0,5039 |
| CD40 | Seasonality_Rainy | -0,1233 | 0,0411 | -3,0016 | 0,0029 | 0,0219 |
| CD5 | Age | -0,0047 | 0,0017 | -2,6736 | 0,0079 | 0,0417 |
| CD5 | Female | 0,0154 | 0,0356 | 0,4344 | 0,6643 | 0,8085 |
| CD5 | BMI | 0,0042 | 0,0039 | 1,0804 | 0,2808 | 0,4901 |
| CD5 | Residency_AreaUrban | 0,0156 | 0,0433 | 0,3596 | 0,7194 | 0,8531 |
| CD5 | Seasonality_Rainy | -0,0286 | 0,0367 | -0,7795 | 0,4362 | 0,6509 |
| CD6 | Age | 0,0014 | 0,0024 | 0,5776 | 0,5639 | 0,7605 |
| CD6 | Female | -0,0736 | 0,0497 | -1,4801 | 0,1399 | 0,3255 |
| CD6 | BMI | 0,0036 | 0,0055 | 0,6617 | 0,5086 | 0,7240 |
| CD6 | Residency_AreaUrban | 0,0486 | 0,0605 | 0,8036 | 0,4223 | 0,6429 |
| CD6 | Seasonality_Rainy | -0,0261 | 0,0513 | -0,5093 | 0,6109 | 0,7767 |
| CD8A | Age | -0,0023 | 0,0050 | -0,4563 | 0,6485 | 0,7954 |
| CD8A | Female | 0,0612 | 0,1016 | 0,6021 | 0,5476 | 0,7531 |
| CD8A | BMI | -0,0162 | 0,0112 | -1,4494 | 0,1482 | 0,3304 |
| CD8A | Residency_AreaUrban | 0,1924 | 0,1237 | 1,5558 | 0,1208 | 0,2993 |
| CD8A | Seasonality_Rainy | -0,0362 | 0,1049 | -0,3446 | 0,7306 | 0,8637 |
| CDCP1 | Age | 0,0285 | 0,0034 | 8,4002 | 0,0000 | 0,0000 |
| CDCP1 | Female | -0,2371 | 0,0693 | -3,4227 | 0,0007 | 0,0067 |
| CDCP1 | BMI | 0,0176 | 0,0076 | 2,3073 | 0,0217 | 0,0863 |
| CDCP1 | Residency_AreaUrban | -0,0066 | 0,0844 | -0,0778 | 0,9381 | 0,9743 |
| CDCP1 | Seasonality_Rainy | -0,1726 | 0,0716 | -2,4118 | 0,0165 | 0,0684 |
| CSF-1 | Age | 0,0004 | 0,0013 | 0,3111 | 0,7559 | 0,8854 |
| CSF-1 | Female | 0,0542 | 0,0259 | 2,0891 | 0,0375 | 0,1285 |
| CSF-1 | BMI | 0,0098 | 0,0029 | 3,4312 | 0,0007 | 0,0066 |
| CSF-1 | Residency_AreaUrban | 0,0090 | 0,0316 | 0,2841 | 0,7766 | 0,8955 |
| CSF-1 | Seasonality_Rainy | -0,0116 | 0,0268 | -0,4312 | 0,6666 | 0,8087 |
| CST5 | Age | 0,0093 | 0,0032 | 2,9078 | 0,0039 | 0,0267 |
| CST5 | Female | -0,2641 | 0,0655 | -4,0298 | 0,0001 | 0,0010 |
| CST5 | BMI | -0,0114 | 0,0072 | -1,5844 | 0,1141 | 0,2892 |
| CST5 | Residency_AreaUrban | -0,0863 | 0,0798 | -1,0808 | 0,2806 | 0,4901 |
| CST5 | Seasonality_Rainy | -0,1205 | 0,0677 | -1,7791 | 0,0762 | 0,2108 |
| CX3CL1 | Age | 0,0026 | 0,0022 | 1,1725 | 0,2419 | 0,4566 |
| CX3CL1 | Female | 0,0283 | 0,0456 | 0,6213 | 0,5348 | 0,7468 |
| CX3CL1 | BMI | -0,0092 | 0,0050 | -1,8422 | 0,0664 | 0,1929 |
| CX3CL1 | Residency_AreaUrban | -0,0810 | 0,0556 | -1,4575 | 0,1460 | 0,3274 |
| CX3CL1 | Seasonality_Rainy | 0,0238 | 0,0471 | 0,5042 | 0,6145 | 0,7767 |
| CXCL1 | Age | -0,0093 | 0,0085 | -1,0903 | 0,2764 | 0,4871 |
| CXCL1 | Female | -0,0879 | 0,1746 | -0,5034 | 0,6151 | 0,7767 |
| CXCL1 | BMI | 0,0239 | 0,0192 | 1,2432 | 0,2147 | 0,4295 |
| CXCL1 | Residency_AreaUrban | -0,2043 | 0,2126 | -0,9610 | 0,3373 | 0,5481 |
| CXCL1 | Seasonality_Rainy | 0,5050 | 0,1803 | 2,8002 | 0,0054 | 0,0314 |
| CXCL10 | Age | 0,0008 | 0,0042 | 0,1875 | 0,8514 | 0,9390 |
| CXCL10 | Female | 0,0228 | 0,0858 | 0,2654 | 0,7909 | 0,9039 |
| CXCL10 | BMI | 0,0189 | 0,0094 | 1,9994 | 0,0464 | 0,1534 |
| CXCL10 | Residency_AreaUrban | -0,0012 | 0,1045 | -0,0111 | 0,9912 | 0,9943 |
| CXCL10 | Seasonality_Rainy | 0,1698 | 0,0887 | 1,9147 | 0,0564 | 0,1726 |
| CXCL11 | Age | 0,0017 | 0,0061 | 0,2834 | 0,7770 | 0,8955 |
| CXCL11 | Female | 0,2232 | 0,1253 | 1,7819 | 0,0757 | 0,2108 |
| CXCL11 | BMI | 0,0100 | 0,0138 | 0,7286 | 0,4668 | 0,6800 |
| CXCL11 | Residency_AreaUrban | 0,0176 | 0,1526 | 0,1155 | 0,9081 | 0,9600 |
| CXCL11 | Seasonality_Rainy | 0,2707 | 0,1294 | 2,0915 | 0,0373 | 0,1285 |
| CXCL5 | Age | -0,0124 | 0,0108 | -1,1512 | 0,2505 | 0,4661 |
| CXCL5 | Female | 0,2685 | 0,2200 | 1,2202 | 0,2233 | 0,4395 |
| CXCL5 | BMI | 0,0466 | 0,0242 | 1,9228 | 0,0554 | 0,1709 |
| CXCL5 | Residency_AreaUrban | -0,5563 | 0,2680 | -2,0759 | 0,0387 | 0,1314 |
| CXCL5 | Seasonality_Rainy | 0,7667 | 0,2273 | 3,3730 | 0,0008 | 0,0076 |
| CXCL6 | Age | -0,0081 | 0,0067 | -1,2108 | 0,2269 | 0,4418 |
| CXCL6 | Female | -0,1400 | 0,1360 | -1,0293 | 0,3041 | 0,5162 |
| CXCL6 | BMI | 0,0169 | 0,0150 | 1,1303 | 0,2592 | 0,4725 |
| CXCL6 | Residency_AreaUrban | -0,2162 | 0,1657 | -1,3050 | 0,1928 | 0,3963 |
| CXCL6 | Seasonality_Rainy | 0,4643 | 0,1405 | 3,3045 | 0,0011 | 0,0094 |
| CXCL9 | Age | 0,0155 | 0,0048 | 3,2124 | 0,0015 | 0,0120 |
| CXCL9 | Female | -0,2947 | 0,0986 | -2,9879 | 0,0030 | 0,0224 |
| CXCL9 | BMI | -0,0062 | 0,0109 | -0,5708 | 0,5685 | 0,7605 |
| CXCL9 | Residency_AreaUrban | 0,0154 | 0,1201 | 0,1285 | 0,8978 | 0,9570 |
| CXCL9 | Seasonality_Rainy | 0,0627 | 0,1019 | 0,6154 | 0,5387 | 0,7482 |
| DNER | Age | 0,0015 | 0,0013 | 1,2154 | 0,2251 | 0,4408 |
| DNER | Female | -0,1608 | 0,0258 | -6,2290 | 0,0000 | 0,0000 |
| DNER | BMI | -0,0055 | 0,0028 | -1,9256 | 0,0551 | 0,1709 |
| DNER | Residency_AreaUrban | -0,0294 | 0,0314 | -0,9363 | 0,3499 | 0,5653 |
| DNER | Seasonality_Rainy | -0,1048 | 0,0267 | -3,9307 | 0,0001 | 0,0013 |
| EN-RAge | Age | -0,0103 | 0,0036 | -2,8670 | 0,0044 | 0,0282 |
| EN-RAge | Female | -0,2638 | 0,0731 | -3,6076 | 0,0004 | 0,0039 |
| EN-RAge | BMI | 0,0216 | 0,0080 | 2,6783 | 0,0078 | 0,0417 |
| EN-RAge | Residency_AreaUrban | 0,0577 | 0,0891 | 0,6474 | 0,5179 | 0,7335 |
| EN-RAge | Seasonality_Rainy | -0,1919 | 0,0755 | -2,5406 | 0,0116 | 0,0562 |
| FGF-19 | Age | 0,0189 | 0,0051 | 3,6715 | 0,0003 | 0,0033 |
| FGF-19 | Female | -0,2616 | 0,1050 | -2,4907 | 0,0133 | 0,0599 |
| FGF-19 | BMI | -0,0194 | 0,0116 | -1,6768 | 0,0946 | 0,2430 |
| FGF-19 | Residency_AreaUrban | 0,0508 | 0,1279 | 0,3968 | 0,6918 | 0,8257 |
| FGF-19 | Seasonality_Rainy | 0,0057 | 0,1085 | 0,0527 | 0,9580 | 0,9819 |
| FGF-21 | Age | 0,0239 | 0,0084 | 2,8518 | 0,0046 | 0,0291 |
| FGF-21 | Female | -0,2658 | 0,1714 | -1,5511 | 0,1219 | 0,2993 |
| FGF-21 | BMI | -0,0182 | 0,0189 | -0,9668 | 0,3344 | 0,5475 |
| FGF-21 | Residency_AreaUrban | -0,2726 | 0,2087 | -1,3063 | 0,1924 | 0,3963 |
| FGF-21 | Seasonality_Rainy | -0,1074 | 0,1770 | -0,6070 | 0,5443 | 0,7515 |
| FGF-23 | Age | 0,0016 | 0,0055 | 0,2870 | 0,7743 | 0,8955 |
| FGF-23 | Female | 0,5069 | 0,1131 | 4,4807 | 0,0000 | 0,0002 |
| FGF-23 | BMI | -0,0013 | 0,0125 | -0,1063 | 0,9154 | 0,9644 |
| FGF-23 | Residency_AreaUrban | 0,1908 | 0,1378 | 1,3847 | 0,1671 | 0,3575 |
| FGF-23 | Seasonality_Rainy | -0,0788 | 0,1169 | -0,6744 | 0,5005 | 0,7178 |
| Flt3L | Age | 0,0086 | 0,0024 | 3,6254 | 0,0003 | 0,0038 |
| Flt3L | Female | 0,1054 | 0,0487 | 2,1626 | 0,0313 | 0,1136 |
| Flt3L | BMI | 0,0064 | 0,0054 | 1,1871 | 0,2361 | 0,4479 |
| Flt3L | Residency_AreaUrban | -0,0166 | 0,0593 | -0,2804 | 0,7794 | 0,8955 |
| Flt3L | Seasonality_Rainy | -0,0270 | 0,0503 | -0,5364 | 0,5921 | 0,7659 |
| HGF | Age | 0,0009 | 0,0019 | 0,4862 | 0,6271 | 0,7852 |
| HGF | Female | -0,0367 | 0,0378 | -0,9696 | 0,3330 | 0,5475 |
| HGF | BMI | 0,0200 | 0,0042 | 4,8000 | 0,0000 | 0,0001 |
| HGF | Residency_AreaUrban | -0,0305 | 0,0461 | -0,6615 | 0,5088 | 0,7240 |
| HGF | Seasonality_Rainy | 0,0662 | 0,0391 | 1,6936 | 0,0913 | 0,2363 |
| IFN-gamma | Age | -0,0054 | 0,0071 | -0,7716 | 0,4409 | 0,6527 |
| IFN-gamma | Female | 0,0768 | 0,1443 | 0,5326 | 0,5947 | 0,7667 |
| IFN-gamma | BMI | 0,0223 | 0,0159 | 1,4069 | 0,1604 | 0,3534 |
| IFN-gamma | Residency_AreaUrban | 0,1390 | 0,1757 | 0,7913 | 0,4294 | 0,6442 |
| IFN-gamma | Seasonality_Rainy | 0,1238 | 0,1490 | 0,8306 | 0,4069 | 0,6272 |
| IL-10RA | Age | -0,0049 | 0,0046 | -1,0700 | 0,2854 | 0,4958 |
| IL-10RA | Female | -0,0216 | 0,0930 | -0,2325 | 0,8163 | 0,9193 |
| IL-10RA | BMI | -0,0018 | 0,0102 | -0,1751 | 0,8611 | 0,9390 |
| IL-10RA | Residency_AreaUrban | 0,0205 | 0,1133 | 0,1811 | 0,8564 | 0,9390 |
| IL-10RA | Seasonality_Rainy | 0,1430 | 0,0961 | 1,4875 | 0,1379 | 0,3229 |
| IL-10RB | Age | 0,0011 | 0,0017 | 0,6331 | 0,5271 | 0,7388 |
| IL-10RB | Female | -0,0999 | 0,0352 | -2,8425 | 0,0048 | 0,0294 |
| IL-10RB | BMI | 0,0067 | 0,0039 | 1,7396 | 0,0829 | 0,2207 |
| IL-10RB | Residency_AreaUrban | -0,0174 | 0,0428 | -0,4064 | 0,6847 | 0,8226 |
| IL-10RB | Seasonality_Rainy | -0,0910 | 0,0363 | -2,5047 | 0,0128 | 0,0584 |
| IL-12B | Age | -0,0037 | 0,0033 | -1,1400 | 0,2552 | 0,4674 |
| IL-12B | Female | 0,3557 | 0,0672 | 5,2958 | 0,0000 | 0,0000 |
| IL-12B | BMI | 0,0155 | 0,0074 | 2,0982 | 0,0367 | 0,1281 |
| IL-12B | Residency_AreaUrban | 0,0806 | 0,0818 | 0,9853 | 0,3253 | 0,5397 |
| IL-12B | Seasonality_Rainy | -0,0512 | 0,0694 | -0,7374 | 0,4614 | 0,6748 |
| IL-15RA | Age | 0,0019 | 0,0015 | 1,3218 | 0,1872 | 0,3891 |
| IL-15RA | Female | -0,0979 | 0,0301 | -3,2508 | 0,0013 | 0,0110 |
| IL-15RA | BMI | 0,0026 | 0,0033 | 0,7959 | 0,4267 | 0,6442 |
| IL-15RA | Residency_AreaUrban | -0,0023 | 0,0367 | -0,0630 | 0,9498 | 0,9789 |
| IL-15RA | Seasonality_Rainy | 0,0012 | 0,0311 | 0,0379 | 0,9698 | 0,9845 |
| IL-17A | Age | 0,0068 | 0,0050 | 1,3697 | 0,1718 | 0,3625 |
| IL-17A | Female | 0,2469 | 0,1015 | 2,4317 | 0,0156 | 0,0671 |
| IL-17A | BMI | -0,0164 | 0,0112 | -1,4672 | 0,1433 | 0,3273 |
| IL-17A | Residency_AreaUrban | -0,0506 | 0,1237 | -0,4089 | 0,6829 | 0,8226 |
| IL-17A | Seasonality_Rainy | -0,0598 | 0,1049 | -0,5696 | 0,5693 | 0,7605 |
| IL-17C | Age | 0,0032 | 0,0030 | 1,0371 | 0,3005 | 0,5123 |
| IL-17C | Female | -0,0690 | 0,0623 | -1,1075 | 0,2689 | 0,4845 |
| IL-17C | BMI | 0,0007 | 0,0069 | 0,0971 | 0,9227 | 0,9644 |
| IL-17C | Residency_AreaUrban | -0,0705 | 0,0759 | -0,9293 | 0,3535 | 0,5686 |
| IL-17C | Seasonality_Rainy | -0,0671 | 0,0644 | -1,0434 | 0,2976 | 0,5097 |
| IL-18R1 | Age | -0,0009 | 0,0022 | -0,4153 | 0,6782 | 0,8201 |
| IL-18R1 | Female | -0,0382 | 0,0452 | -0,8455 | 0,3985 | 0,6169 |
| IL-18R1 | BMI | 0,0217 | 0,0050 | 4,3724 | 0,0000 | 0,0003 |
| IL-18R1 | Residency_AreaUrban | 0,0014 | 0,0550 | 0,0254 | 0,9798 | 0,9905 |
| IL-18R1 | Seasonality_Rainy | -0,0689 | 0,0467 | -1,4768 | 0,1407 | 0,3255 |
| IL10 | Age | 0,0086 | 0,0046 | 1,8718 | 0,0622 | 0,1840 |
| IL10 | Female | -0,1605 | 0,0934 | -1,7185 | 0,0867 | 0,2275 |
| IL10 | BMI | -0,0056 | 0,0103 | -0,5415 | 0,5886 | 0,7641 |
| IL10 | Residency_AreaUrban | 0,1248 | 0,1137 | 1,0971 | 0,2734 | 0,4863 |
| IL10 | Seasonality_Rainy | 0,0300 | 0,0965 | 0,3107 | 0,7562 | 0,8854 |
| IL18 | Age | 0,0045 | 0,0037 | 1,2065 | 0,2285 | 0,4427 |
| IL18 | Female | -0,3011 | 0,0756 | -3,9832 | 0,0001 | 0,0012 |
| IL18 | BMI | 0,0179 | 0,0083 | 2,1557 | 0,0319 | 0,1136 |
| IL18 | Residency_AreaUrban | -0,0710 | 0,0921 | -0,7715 | 0,4410 | 0,6527 |
| IL18 | Seasonality_Rainy | -0,1734 | 0,0781 | -2,2200 | 0,0271 | 0,1014 |
| IL6 | Age | 0,0056 | 0,0051 | 1,1056 | 0,2698 | 0,4845 |
| IL6 | Female | 0,4481 | 0,1038 | 4,3173 | 0,0000 | 0,0004 |
| IL6 | BMI | 0,0403 | 0,0114 | 3,5241 | 0,0005 | 0,0050 |
| IL6 | Residency_AreaUrban | -0,0381 | 0,1264 | -0,3011 | 0,7636 | 0,8894 |
| IL6 | Seasonality_Rainy | 0,0551 | 0,1072 | 0,5141 | 0,6075 | 0,7767 |
| IL7 | Age | -0,0020 | 0,0044 | -0,4553 | 0,6492 | 0,7954 |
| IL7 | Female | -0,0650 | 0,0908 | -0,7159 | 0,4746 | 0,6859 |
| IL7 | BMI | 0,0115 | 0,0100 | 1,1473 | 0,2521 | 0,4664 |
| IL7 | Residency_AreaUrban | 0,0903 | 0,1105 | 0,8166 | 0,4148 | 0,6368 |
| IL7 | Seasonality_Rainy | 0,0066 | 0,0937 | 0,0702 | 0,9441 | 0,9757 |
| IL8 | Age | 0,0007 | 0,0061 | 0,1228 | 0,9024 | 0,9570 |
| IL8 | Female | -0,1259 | 0,1237 | -1,0182 | 0,3094 | 0,5196 |
| IL8 | BMI | 0,0025 | 0,0136 | 0,1802 | 0,8571 | 0,9390 |
| IL8 | Residency_AreaUrban | -0,2836 | 0,1506 | -1,8833 | 0,0606 | 0,1808 |
| IL8 | Seasonality_Rainy | 0,1179 | 0,1277 | 0,9227 | 0,3569 | 0,5716 |
| LAP.TGF-beta-1 | Age | 0,0011 | 0,0026 | 0,4405 | 0,6599 | 0,8058 |
| LAP.TGF-beta-1 | Female | 0,0078 | 0,0521 | 0,1489 | 0,8817 | 0,9487 |
| LAP.TGF-beta-1 | BMI | 0,0068 | 0,0057 | 1,1921 | 0,2341 | 0,4465 |
| LAP.TGF-beta-1 | Residency_AreaUrban | -0,0372 | 0,0635 | -0,5858 | 0,5584 | 0,7605 |
| LAP.TGF-beta-1 | Seasonality_Rainy | 0,0701 | 0,0538 | 1,3020 | 0,1939 | 0,3963 |
| LIF-R | Age | 0,0027 | 0,0014 | 2,0277 | 0,0434 | 0,1461 |
| LIF-R | Female | -0,0244 | 0,0277 | -0,8836 | 0,3776 | 0,5927 |
| LIF-R | BMI | -0,0084 | 0,0030 | -2,7507 | 0,0063 | 0,0353 |
| LIF-R | Residency_AreaUrban | -0,0285 | 0,0337 | -0,8461 | 0,3981 | 0,6169 |
| LIF-R | Seasonality_Rainy | -0,0693 | 0,0286 | -2,4268 | 0,0158 | 0,0672 |
| MCP-1 | Age | 0,0080 | 0,0030 | 2,6224 | 0,0092 | 0,0471 |
| MCP-1 | Female | -0,1336 | 0,0620 | -2,1550 | 0,0319 | 0,1136 |
| MCP-1 | BMI | 0,0161 | 0,0068 | 2,3651 | 0,0186 | 0,0750 |
| MCP-1 | Residency_AreaUrban | -0,0174 | 0,0755 | -0,2310 | 0,8174 | 0,9193 |
| MCP-1 | Seasonality_Rainy | 0,3085 | 0,0641 | 4,8153 | 0,0000 | 0,0001 |
| MCP-2 | Age | 0,0020 | 0,0035 | 0,5716 | 0,5680 | 0,7605 |
| MCP-2 | Female | -0,1400 | 0,0713 | -1,9639 | 0,0504 | 0,1622 |
| MCP-2 | BMI | 0,0282 | 0,0078 | 3,5886 | 0,0004 | 0,0041 |
| MCP-2 | Residency_AreaUrban | -0,0223 | 0,0868 | -0,2570 | 0,7974 | 0,9071 |
| MCP-2 | Seasonality_Rainy | 0,1685 | 0,0736 | 2,2881 | 0,0228 | 0,0897 |
| MCP-3 | Age | 0,0062 | 0,0035 | 1,7708 | 0,0776 | 0,2108 |
| MCP-3 | Female | -0,0565 | 0,0715 | -0,7902 | 0,4300 | 0,6442 |
| MCP-3 | BMI | 0,0298 | 0,0079 | 3,7882 | 0,0002 | 0,0022 |
| MCP-3 | Residency_AreaUrban | 0,0975 | 0,0871 | 1,1194 | 0,2638 | 0,4785 |
| MCP-3 | Seasonality_Rainy | -0,0152 | 0,0739 | -0,2059 | 0,8370 | 0,9385 |
| MCP-4 | Age | 0,0091 | 0,0046 | 1,9544 | 0,0515 | 0,1644 |
| MCP-4 | Female | 0,0169 | 0,0949 | 0,1783 | 0,8586 | 0,9390 |
| MCP-4 | BMI | 0,0266 | 0,0104 | 2,5476 | 0,0113 | 0,0559 |
| MCP-4 | Residency_AreaUrban | 0,0463 | 0,1156 | 0,4002 | 0,6893 | 0,8254 |
| MCP-4 | Seasonality_Rainy | 0,5117 | 0,0981 | 5,2188 | 0,0000 | 0,0000 |
| MMP-1 | Age | 0,0113 | 0,0064 | 1,7701 | 0,0777 | 0,2108 |
| MMP-1 | Female | -0,3555 | 0,1309 | -2,7153 | 0,0070 | 0,0386 |
| MMP-1 | BMI | 0,0253 | 0,0144 | 1,7572 | 0,0799 | 0,2141 |
| MMP-1 | Residency_AreaUrban | 0,0195 | 0,1595 | 0,1224 | 0,9026 | 0,9570 |
| MMP-1 | Seasonality_Rainy | 0,3948 | 0,1353 | 2,9190 | 0,0038 | 0,0267 |
| MMP-10 | Age | 0,0104 | 0,0037 | 2,8036 | 0,0054 | 0,0314 |
| MMP-10 | Female | 0,0117 | 0,0757 | 0,1546 | 0,8772 | 0,9487 |
| MMP-10 | BMI | -0,0247 | 0,0083 | -2,9664 | 0,0032 | 0,0235 |
| MMP-10 | Residency_AreaUrban | 0,0235 | 0,0922 | 0,2546 | 0,7992 | 0,9071 |
| MMP-10 | Seasonality_Rainy | -0,1383 | 0,0782 | -1,7678 | 0,0781 | 0,2108 |
| NT-3 | Age | -0,0024 | 0,0028 | -0,8508 | 0,3955 | 0,6169 |
| NT-3 | Female | -0,0565 | 0,0582 | -0,9710 | 0,3323 | 0,5475 |
| NT-3 | BMI | -0,0065 | 0,0064 | -1,0206 | 0,3082 | 0,5196 |
| NT-3 | Residency_AreaUrban | 0,0123 | 0,0709 | 0,1729 | 0,8629 | 0,9390 |
| NT-3 | Seasonality_Rainy | -0,0696 | 0,0601 | -1,1577 | 0,2479 | 0,4655 |
| OPG | Age | 0,0033 | 0,0027 | 1,2266 | 0,2209 | 0,4371 |
| OPG | Female | 0,0655 | 0,0548 | 1,1951 | 0,2329 | 0,4465 |
| OPG | BMI | -0,0003 | 0,0060 | -0,0555 | 0,9558 | 0,9819 |
| OPG | Residency_AreaUrban | -0,1601 | 0,0667 | -2,3982 | 0,0171 | 0,0701 |
| OPG | Seasonality_Rainy | 0,0620 | 0,0566 | 1,0942 | 0,2747 | 0,4863 |
| OSM | Age | -0,0131 | 0,0046 | -2,8258 | 0,0050 | 0,0304 |
| OSM | Female | -0,0609 | 0,0949 | -0,6419 | 0,5214 | 0,7335 |
| OSM | BMI | 0,0316 | 0,0104 | 3,0233 | 0,0027 | 0,0209 |
| OSM | Residency_AreaUrban | -0,0230 | 0,1156 | -0,1989 | 0,8425 | 0,9390 |
| OSM | Seasonality_Rainy | 0,1431 | 0,0981 | 1,4590 | 0,1456 | 0,3274 |
| PD-L1 | Age | -0,0024 | 0,0021 | -1,1508 | 0,2507 | 0,4661 |
| PD-L1 | Female | -0,1755 | 0,0433 | -4,0532 | 0,0001 | 0,0009 |
| PD-L1 | BMI | 0,0028 | 0,0048 | 0,5913 | 0,5547 | 0,7602 |
| PD-L1 | Residency_AreaUrban | -0,0466 | 0,0527 | -0,8827 | 0,3781 | 0,5927 |
| PD-L1 | Seasonality_Rainy | -0,0808 | 0,0447 | -1,8055 | 0,0720 | 0,2048 |
| SCF | Age | -0,0019 | 0,0019 | -1,0161 | 0,3104 | 0,5196 |
| SCF | Female | 0,1305 | 0,0386 | 3,3764 | 0,0008 | 0,0076 |
| SCF | BMI | -0,0129 | 0,0043 | -3,0420 | 0,0025 | 0,0201 |
| SCF | Residency_AreaUrban | -0,0888 | 0,0471 | -1,8867 | 0,0601 | 0,1808 |
| SCF | Seasonality_Rainy | -0,0594 | 0,0399 | -1,4890 | 0,1375 | 0,3229 |
| SIRT2 | Age | -0,0154 | 0,0069 | -2,2255 | 0,0268 | 0,1010 |
| SIRT2 | Female | -0,3578 | 0,1418 | -2,5241 | 0,0121 | 0,0574 |
| SIRT2 | BMI | 0,0385 | 0,0156 | 2,4665 | 0,0142 | 0,0632 |
| SIRT2 | Residency_AreaUrban | -0,0948 | 0,1727 | -0,5489 | 0,5835 | 0,7641 |
| SIRT2 | Seasonality_Rainy | 0,1293 | 0,1465 | 0,8830 | 0,3779 | 0,5927 |
| SLAMF1 | Age | 0,0071 | 0,0017 | 4,1522 | 0,0000 | 0,0007 |
| SLAMF1 | Female | -0,0880 | 0,0348 | -2,5278 | 0,0120 | 0,0574 |
| SLAMF1 | BMI | -0,0053 | 0,0038 | -1,3847 | 0,1671 | 0,3575 |
| SLAMF1 | Residency_AreaUrban | 0,0452 | 0,0424 | 1,0656 | 0,2874 | 0,4970 |
| SLAMF1 | Seasonality_Rainy | 0,0013 | 0,0360 | 0,0361 | 0,9712 | 0,9845 |
| ST1A1 | Age | -0,0136 | 0,0069 | -1,9771 | 0,0489 | 0,1588 |
| ST1A1 | Female | -0,3389 | 0,1401 | -2,4198 | 0,0161 | 0,0677 |
| ST1A1 | BMI | 0,0284 | 0,0154 | 1,8399 | 0,0667 | 0,1929 |
| ST1A1 | Residency_AreaUrban | -0,1876 | 0,1706 | -1,0999 | 0,2722 | 0,4863 |
| ST1A1 | Seasonality_Rainy | 0,0383 | 0,1447 | 0,2646 | 0,7915 | 0,9039 |
| STAMBP | Age | -0,0102 | 0,0048 | -2,1209 | 0,0347 | 0,1223 |
| STAMBP | Female | -0,4075 | 0,0980 | -4,1579 | 0,0000 | 0,0007 |
| STAMBP | BMI | 0,0243 | 0,0108 | 2,2493 | 0,0252 | 0,0971 |
| STAMBP | Residency_AreaUrban | -0,0945 | 0,1194 | -0,7921 | 0,4289 | 0,6442 |
| STAMBP | Seasonality_Rainy | 0,0817 | 0,1012 | 0,8068 | 0,4204 | 0,6427 |
| TGF-alpha | Age | -0,0021 | 0,0022 | -0,9601 | 0,3377 | 0,5481 |
| TGF-alpha | Female | -0,0647 | 0,0440 | -1,4698 | 0,1426 | 0,3273 |
| TGF-alpha | BMI | 0,0076 | 0,0048 | 1,5753 | 0,1162 | 0,2905 |
| TGF-alpha | Residency_AreaUrban | 0,0172 | 0,0536 | 0,3209 | 0,7485 | 0,8820 |
| TGF-alpha | Seasonality_Rainy | 0,0449 | 0,0455 | 0,9869 | 0,3245 | 0,5397 |
| TNF | Age | -0,0004 | 0,0043 | -0,0973 | 0,9226 | 0,9644 |
| TNF | Female | -0,0418 | 0,0879 | -0,4755 | 0,6348 | 0,7882 |
| TNF | BMI | 0,0088 | 0,0097 | 0,9072 | 0,3650 | 0,5821 |
| TNF | Residency_AreaUrban | -0,1320 | 0,1071 | -1,2325 | 0,2187 | 0,4351 |
| TNF | Seasonality_Rainy | 0,0438 | 0,0908 | 0,4824 | 0,6298 | 0,7852 |
| TNFB | Age | -0,0087 | 0,0021 | -4,2230 | 0,0000 | 0,0006 |
| TNFB | Female | 0,2140 | 0,0420 | 5,1003 | 0,0000 | 0,0000 |
| TNFB | BMI | 0,0069 | 0,0046 | 1,5037 | 0,1337 | 0,3170 |
| TNFB | Residency_AreaUrban | -0,0745 | 0,0511 | -1,4584 | 0,1457 | 0,3274 |
| TNFB | Seasonality_Rainy | -0,0672 | 0,0433 | -1,5501 | 0,1221 | 0,2993 |
| TNFRSF9 | Age | -0,0034 | 0,0024 | -1,4342 | 0,1525 | 0,3379 |
| TNFRSF9 | Female | -0,0943 | 0,0487 | -1,9355 | 0,0538 | 0,1703 |
| TNFRSF9 | BMI | 0,0004 | 0,0054 | 0,0752 | 0,9401 | 0,9743 |
| TNFRSF9 | Residency_AreaUrban | -0,0766 | 0,0593 | -1,2912 | 0,1976 | 0,4017 |
| TNFRSF9 | Seasonality_Rainy | -0,0828 | 0,0503 | -1,6456 | 0,1009 | 0,2573 |
| TNFSF14 | Age | -0,0047 | 0,0028 | -1,7100 | 0,0883 | 0,2300 |
| TNFSF14 | Female | -0,0364 | 0,0567 | -0,6421 | 0,5213 | 0,7335 |
| TNFSF14 | BMI | 0,0282 | 0,0062 | 4,5176 | 0,0000 | 0,0002 |
| TNFSF14 | Residency_AreaUrban | -0,0098 | 0,0691 | -0,1415 | 0,8875 | 0,9491 |
| TNFSF14 | Seasonality_Rainy | 0,0319 | 0,0586 | 0,5451 | 0,5861 | 0,7641 |
| TRAIL | Age | -0,0047 | 0,0016 | -2,8672 | 0,0044 | 0,0282 |
| TRAIL | Female | -0,1547 | 0,0334 | -4,6272 | 0,0000 | 0,0001 |
| TRAIL | BMI | 0,0083 | 0,0037 | 2,2536 | 0,0249 | 0,0970 |
| TRAIL | Residency_AreaUrban | -0,0221 | 0,0407 | -0,5424 | 0,5879 | 0,7641 |
| TRAIL | Seasonality_Rainy | -0,0050 | 0,0345 | -0,1453 | 0,8846 | 0,9487 |
| TRANCE | Age | -0,0185 | 0,0036 | -5,0800 | 0,0000 | 0,0000 |
| TRANCE | Female | -0,0457 | 0,0745 | -0,6136 | 0,5400 | 0,7482 |
| TRANCE | BMI | 0,0265 | 0,0082 | 3,2283 | 0,0014 | 0,0116 |
| TRANCE | Residency_AreaUrban | 0,0415 | 0,0907 | 0,4576 | 0,6476 | 0,7954 |
| TRANCE | Seasonality_Rainy | -0,2216 | 0,0769 | -2,8808 | 0,0042 | 0,0282 |
| TWEAK | Age | -0,0048 | 0,0019 | -2,5039 | 0,0128 | 0,0584 |
| TWEAK | Female | -0,0843 | 0,0391 | -2,1566 | 0,0318 | 0,1136 |
| TWEAK | BMI | 0,0008 | 0,0043 | 0,1837 | 0,8543 | 0,9390 |
| TWEAK | Residency_AreaUrban | -0,0850 | 0,0476 | -1,7840 | 0,0754 | 0,2108 |
| TWEAK | Seasonality_Rainy | -0,0204 | 0,0404 | -0,5038 | 0,6148 | 0,7767 |
| uPA | Age | -0,0047 | 0,0027 | -1,7723 | 0,0773 | 0,2108 |
| uPA | Female | -0,0761 | 0,0545 | -1,3960 | 0,1637 | 0,3571 |
| uPA | BMI | 0,0076 | 0,0060 | 1,2719 | 0,2044 | 0,4109 |
| uPA | Residency_AreaUrban | -0,1732 | 0,0664 | -2,6079 | 0,0095 | 0,0477 |
| uPA | Seasonality_Rainy | -0,0058 | 0,0563 | -0,1023 | 0,9186 | 0,9644 |
| VEGFA | Age | 0,0036 | 0,0018 | 2,0031 | 0,0460 | 0,1534 |
| VEGFA | Female | 0,0004 | 0,0363 | 0,0105 | 0,9917 | 0,9943 |
| VEGFA | BMI | 0,0153 | 0,0040 | 3,8210 | 0,0002 | 0,0020 |
| VEGFA | Residency_AreaUrban | -0,0610 | 0,0442 | -1,3804 | 0,1684 | 0,3582 |
| VEGFA | Seasonality_Rainy | 0,0112 | 0,0375 | 0,2999 | 0,7644 | 0,8894 |

**Table S3.** Comparison inflammatory clusters analysis

| Parameters |  | Lower expression<br>inflammatory cluster<br>(N = 167) | Higher expression<br>inflammatory cluster<br>(N = 124) |  | Statistics |
| --- | --- | --- | --- | --- | --- |
| age | Median | 29,63 | 32,75 | 0.010605 | Mann-Whitney |
|  | IQR | 16,52 | 17,14 |  |  |
|  | Quantile1 | 18,13 | 17,76 |  |  |
|  | Quantile2 | 22,25 | 25,24 |  |  |
|  | Quantile3 | 29,63 | 32,75 |  |  |
|  | Quantile4 | 38,77 | 42,38 |  |  |
|  | Quantile5 | 66,43 | 59,53 |  |  |
| BMI | Median | 23,45 | 24,40 | 0.1362899 | Mann-Whitney |
|  | IQR | 5,50 | 6,89 |  |  |
|  | Quantile1 | 16,41 | 17,04 |  |  |
|  | Quantile2 | 21,33 | 21,79 |  |  |
|  | Quantile3 | 23,45 | 24,40 |  |  |
|  | Quantile4 | 26,83 | 28,69 |  |  |
|  | Quantile5 | 41,91 | 39,84 |  |  |
| Gender | Female | 89/167 (53.3%) | 57/124 (46.0%) | 0.2638188 | Chi-square |
|  | Male | 78/167 (46.7%) | 67/124 (54.0%) |  |  |
| Residency_Area | Rural | 33/167 (19.8%) | 30/124 (24.2%) | 0.4448202 | Chi-square |
|  | Urban | 134/167 (80.2%) | 94/124 (75.8%) |  |  |
| Seasonality | Dry | 127/167 (76.0%) | 85/124 (68.5%) | 0.1973038 | Chi-square |
|  | Rainy | 40/167 (24.0%) | 39/124 (31.5%) |  |  |
| Food Cluster | Healthier food cluster | 89/167 (53.3%) | 46/124 (37.1%) | 0.008769825 | Chi-square |
|  | Less healthy food cluster | 78/167 (46.7%) | 78/124 (62.9%) | 0.0005506364 | Log regression model corrected for age |

**Table S4.**
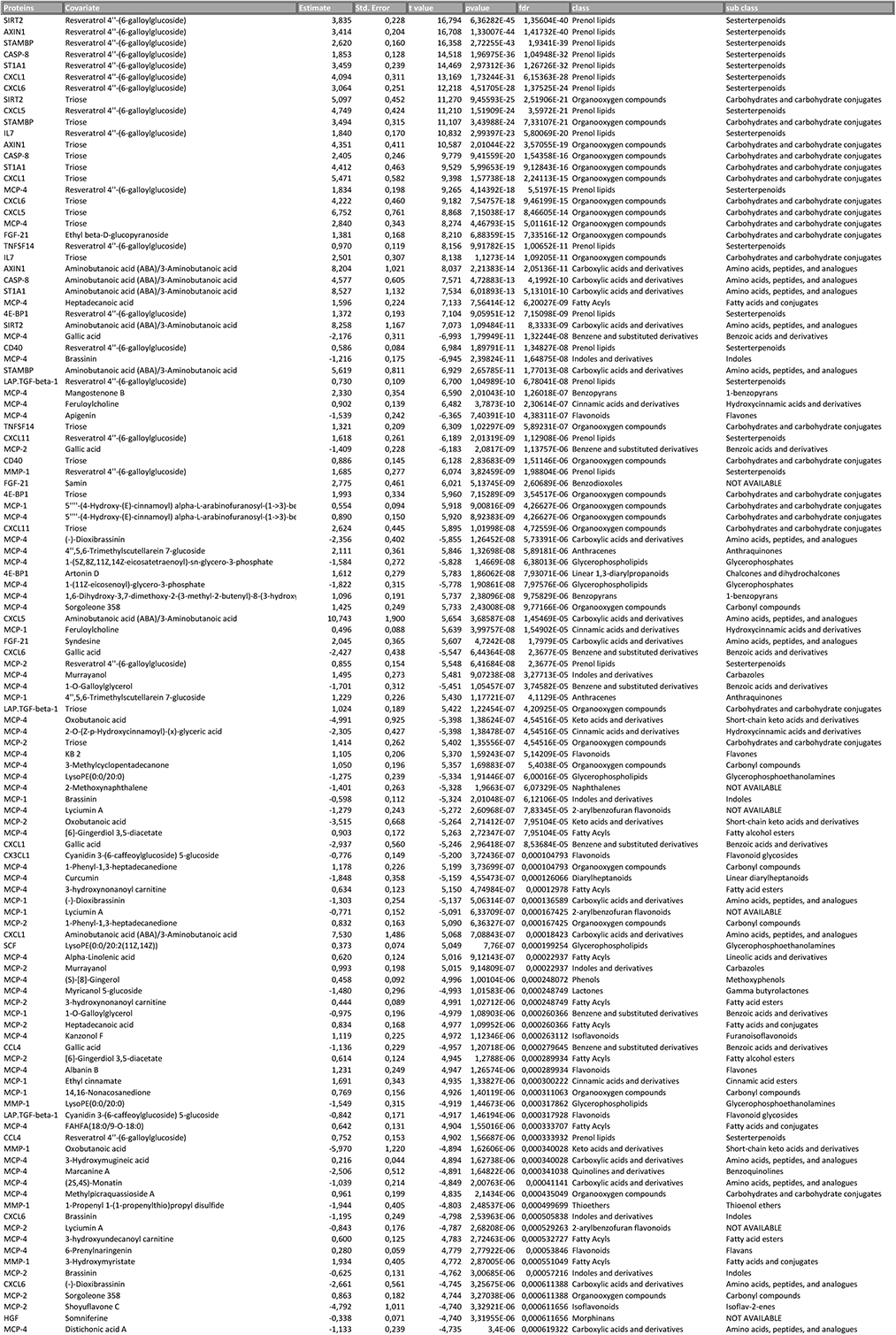
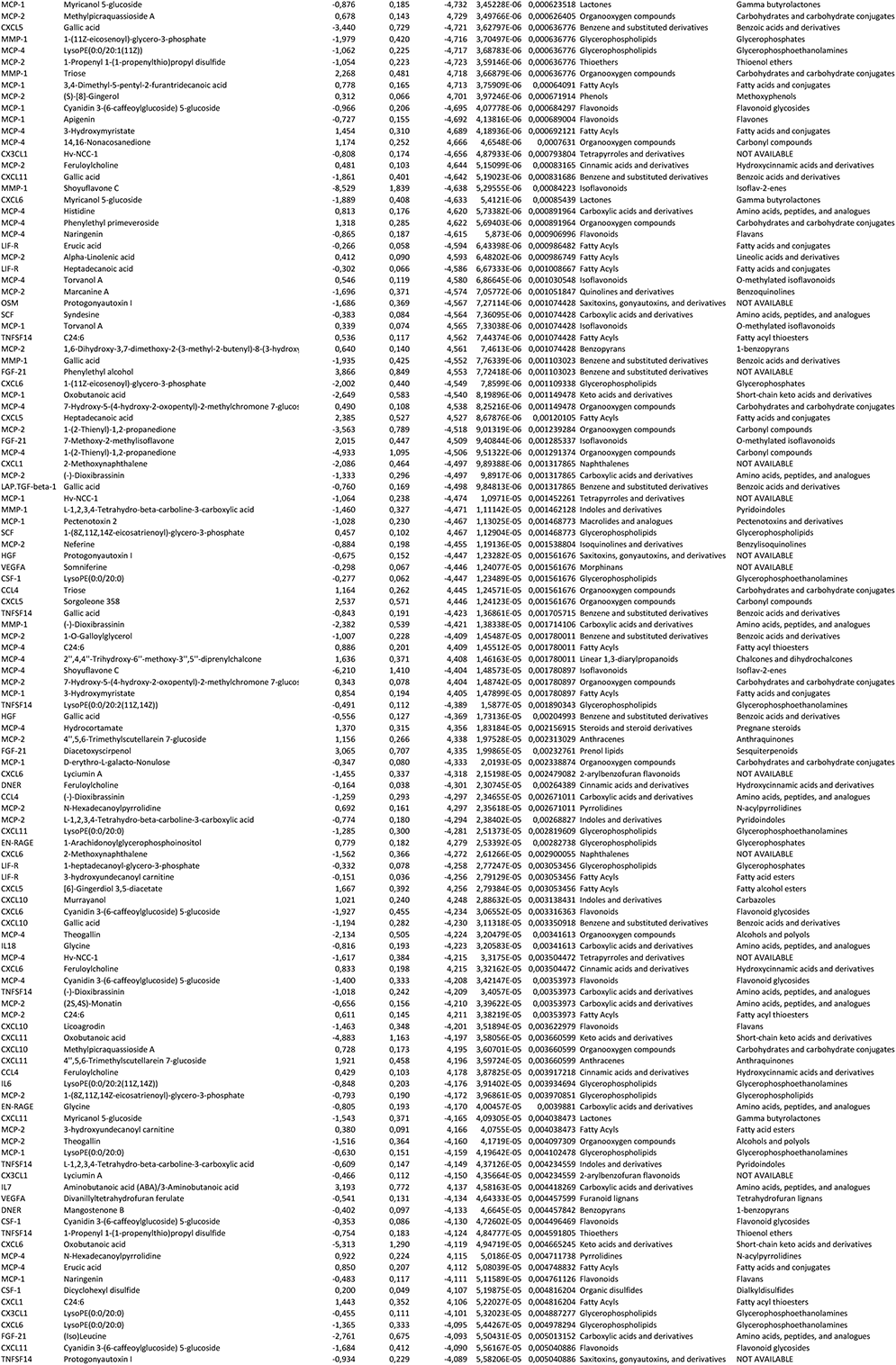
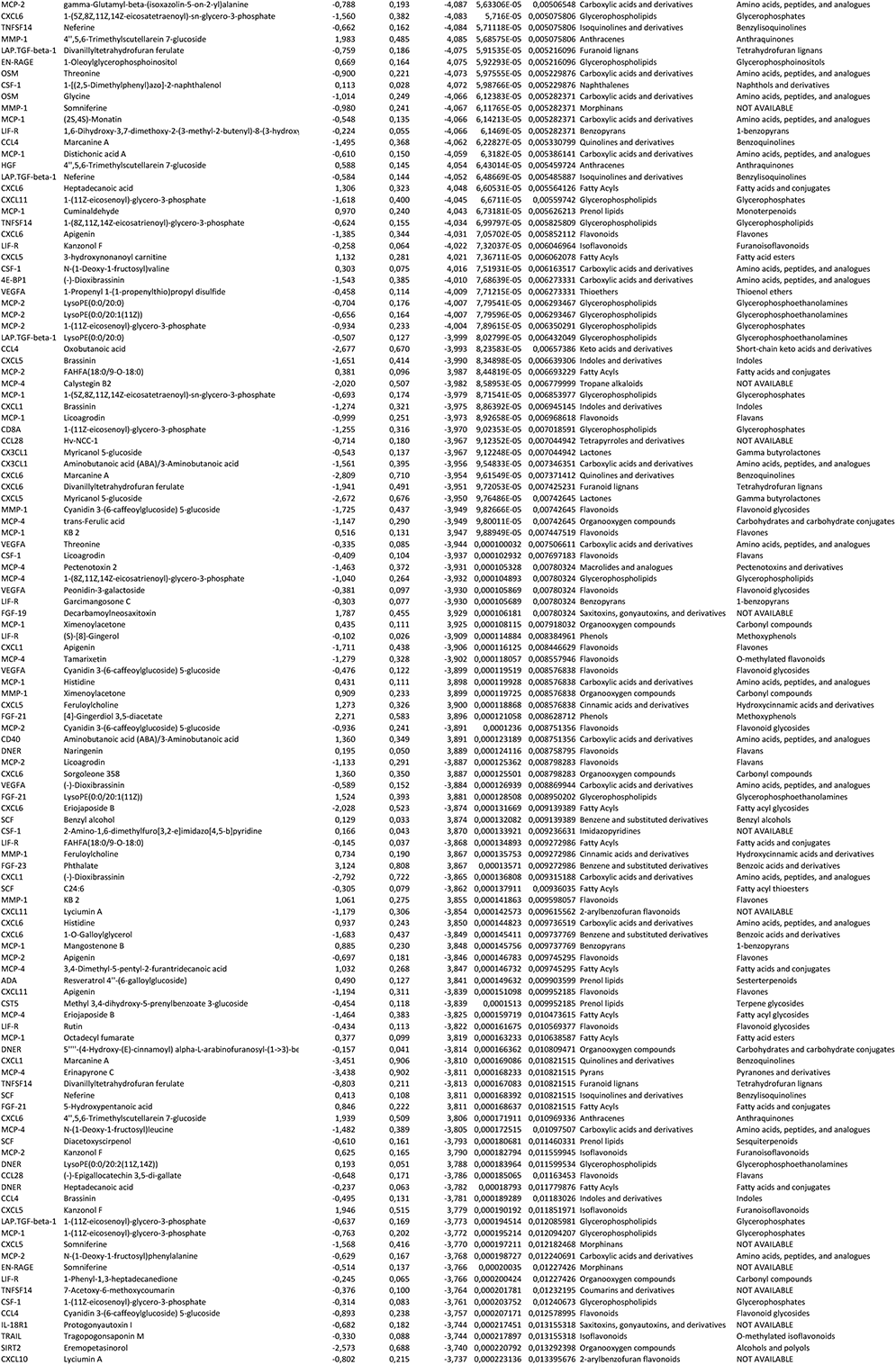
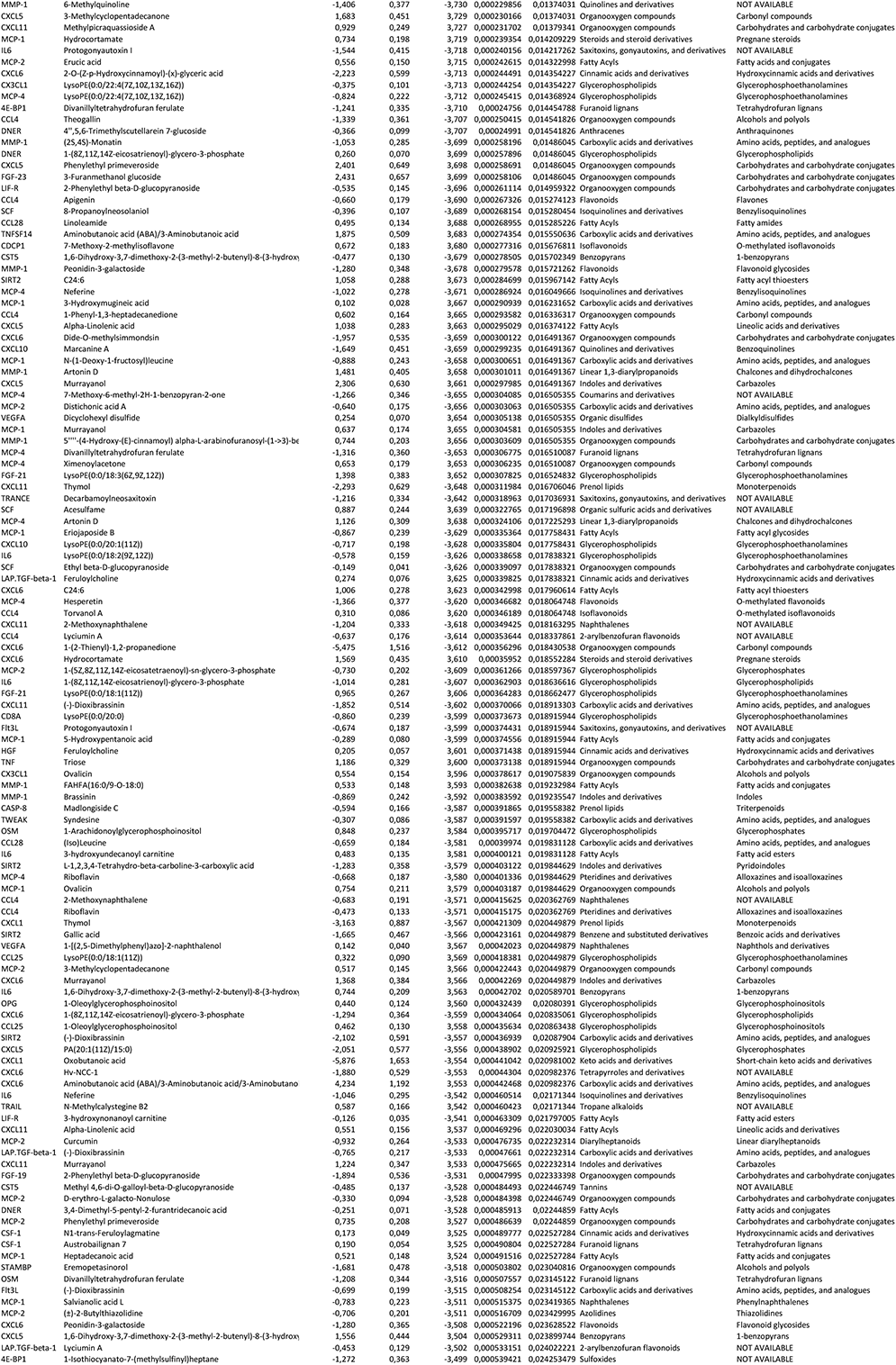
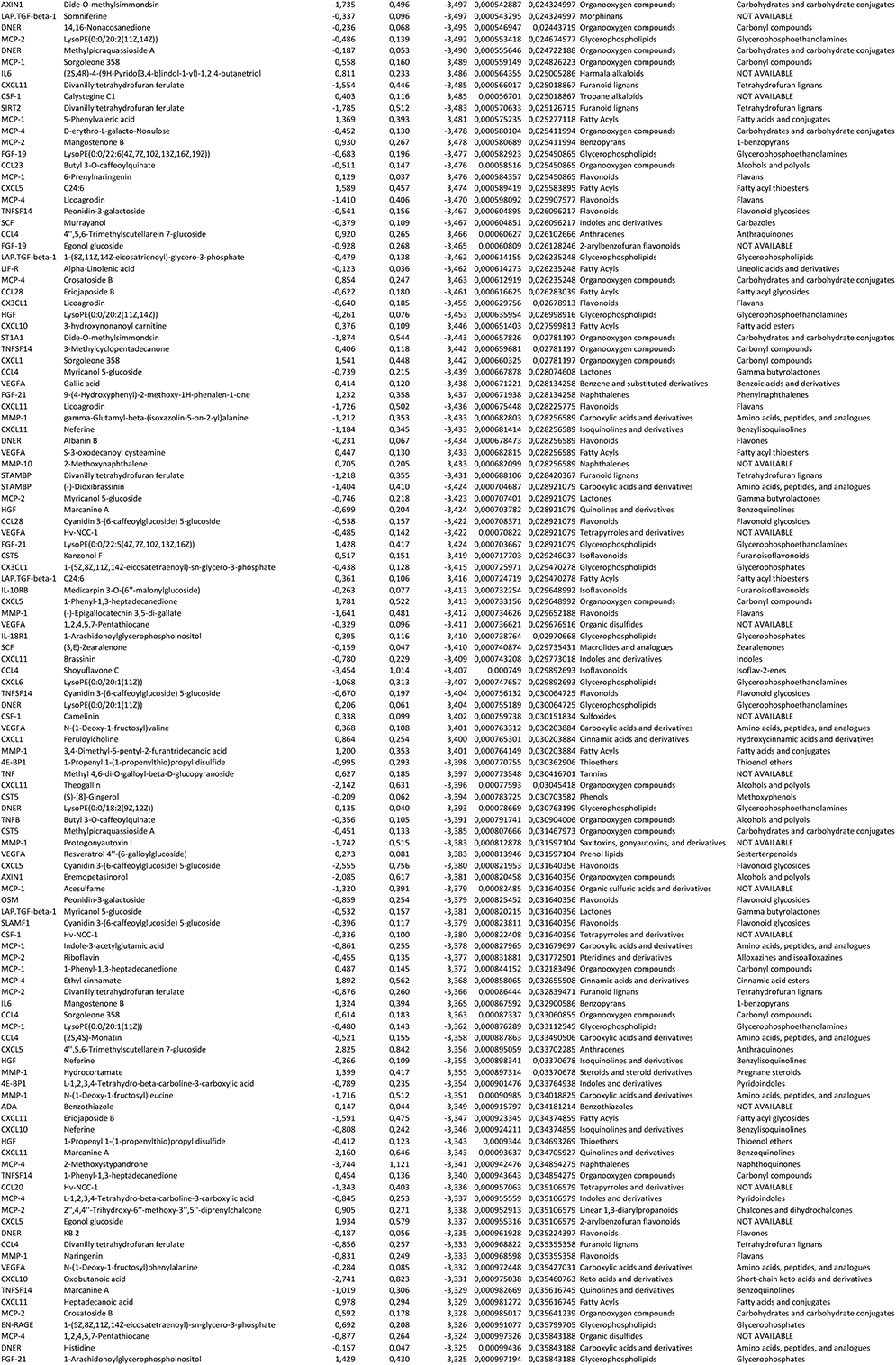
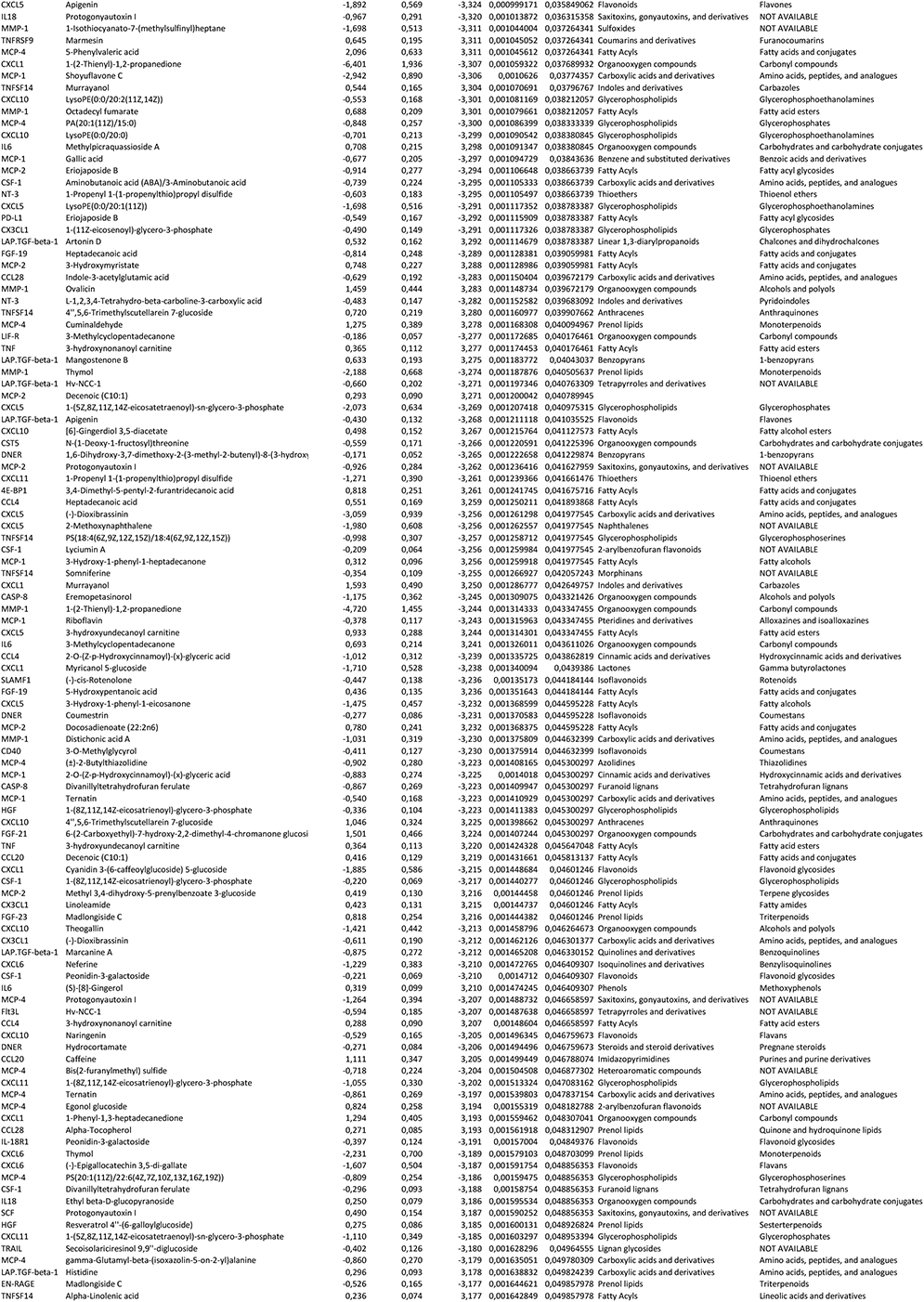
Multivariate linear regression analysis between plasma metabolites and inflammation-related proteins in Tanzanian cohort

## References

Abbas, Z., Lutale, J., & Ahren, B. (2004). Leptin levels in obese and non-obese african and caucasian subjects with type 2 diabetes.

Assarsson, E., Lundberg, M., Holmquist, G., Björkesten, J., Bucht Thorsen, S., Ekman, D., … Edfeldt, G. (2014). Homogenous 96-plex PEA immunoassay exhibiting high sensitivity, specificity, and excellent scalability. PLoS One, 9(4), e95192.

Beaglehole, R., Bonita, R., Horton, R., Adams, C., Alleyne, G., Asaria, P., … Casswell, S. (2011). Priority actions for the non-communicable disease crisis. The lancet, 377(9775), 1438–1447.

Bickler, S. W., Wang, A., Amin, S., Halbach, J., Lizardo, R., Cauvi, D. M., & De Maio, A. (2018). Urbanization in sub-Saharan Africa: declining rates of chronic and recurrent infection and their possible role in the origins of non-communicable diseases. World journal of surgery, 42(6), 1617–1628.

Bielinski, S. J., Berardi, C., Decker, P. A., Larson, N. B., Bell, E. J., Pankow, J. S., … Wassel, C. L. (2017). Hepatocyte growth factor demonstrates racial heterogeneity as a biomarker for coronary heart disease. Heart, 103(15), 1185–1193.

Boahen, C. K., Temba, G. S., Kullaya, V. I., Matzaraki, V., Joosten, L. A., Kibiki, G., … Netea, M. G. (2022). A functional genomics approach in Tanzanian population identifies distinct genetic regulators of cytokine production compared to European population. The American Journal of Human Genetics.

Brodin, P., & Davis, M. M. (2017). Human immune system variation. Nature reviews immunology, 17(1), 21–29.

Choi, Y. J., Kim, N., Chang, H., Lee, H. S., Park, S. M., Park, J. H., … Lee, D. H. (2015). Helicobacter pylori-induced epithelial-mesenchymal transition, a potential role of gastric cancer initiation and an emergence of stem cells. Carcinogenesis, 36(5), 553–563.

Das, B., Das, M., Kalita, A., & Baro, M. R. (2021). The role of Wnt pathway in obesity induced inflammation and diabetes: a review. Journal of Diabetes & Metabolic Disorders, 20(2), 1871–1882.

Enroth, S., Hallmans, G., Grankvist, K., & Gyllensten, U. (2016). Effects of Long-Term Storage Time and Original Sampling Month on Biobank Plasma Protein Concentrations. EBioMedicine, 12, 309–314. doi:10.1016/j.ebiom.2016.08.038

Fuhrer, T., Heer, D., Begemann, B., & Zamboni, N. (2011). High-throughput, accurate mass metabolome profiling of cellular extracts by flow injection-time-of-flight mass spectrometry. Anal Chem, 83(18), 7074–7080. doi:10.1021/ac201267k

Garcia, D. L., Bracci, P. M., Guevarra, D. M., & Sieffert, N. (2014). International Society for Biological and Environmental Repositories (ISBER) tools for the biobanking community. Biopreservation and Biobanking, 12(6), 435–436.

Guarner, V., & Rubio-Ruiz, M. (2015). Aging and Health-A Systems Biology Perspective. Interdiscipl Top Gerontol Basel, Karger, 40, 99–106.

He, Y., Davies, C. M., Harrington, B. S., Hellmers, L., Sheng, Y., Broomfield, A., … Wu, A. (2020). CDCP1 enhances Wnt signaling in colorectal cancer promoting nuclear localization of β-catenin and E-cadherin. Oncogene, 39(1), 219–233.

Hemler, E. C., & Hu, F. B. (2019). Plant-Based Diets for Cardiovascular Disease Prevention: All Plant Foods Are Not Created Equal. Curr Atheroscler Rep, 21(5), 18. doi:10.1007/s11883-019-0779-5

Hill, C. M., Berthoud, H.-R., Münzberg, H., & Morrison, C. D. (2018). Homeostatic sensing of dietary protein restriction: a case for FGF21. Frontiers in neuroendocrinology, 51, 125–131.

Jridi, I., Canté-Barrett, K., Pike-Overzet, K., & Staal, F. J. (2021). Inflammation and Wnt signaling: target for immunomodulatory therapy? Frontiers in Cell and Developmental Biology, 1854.

Karlsson, E. K., Kwiatkowski, D. P., & Sabeti, P. C. (2014). Natural selection and infectious disease in human populations. Nature Reviews Genetics, 15(6), 379–393.

Kikuchi, A. (1999). Roles of Axin in the Wnt signalling pathway. Cellular signalling, 11(11), 777–788.

Kim, H., Castellon-Chicas, M. J., Arbizu, S., Talcott, S. T., Drury, N. L., Smith, S., & Mertens-Talcott, S. U. (2021). Mango (Mangifera indica L.) Polyphenols: Anti-Inflammatory Intestinal Microbial Health Benefits, and Associated Mechanisms of Actions. Molecules, 26(9), 2732.

Lee, J. E., Kim, S. Y., & Shin, S. Y. (2015). Effect of Repeated Freezing and Thawing on Biomarker Stability in Plasma and Serum Samples. Osong Public Health Res Perspect, 6(6), 357–362. doi:10.1016/j.phrp.2015.11.005

Liberale, L., Montecucco, F., Tardif, J.-C., Libby, P., & Camici, G. G. (2020). Inflamm-ageing: the role of inflammation in age-dependent cardiovascular disease. European Heart Journal, 41(31), 2974–2982.

Liston, A., Humblet-Baron, S., Duffy, D., & Goris, A. (2021). Human immune diversity: from evolution to modernity. Nature immunology, 22(12), 1479–1489.

Lundsgaard, A.-M., Fritzen, A. M., Sjøberg, K. A., Myrmel, L. S., Madsen, L., Wojtaszewski, J. F., … Kiens, B. (2017). Circulating FGF21 in humans is potently induced by short term overfeeding of carbohydrates. Molecular metabolism, 6(1), 22–29.

Ma, B., & Hottiger, M. O. (2016). Crosstalk between Wnt/β-catenin and NF-κB signaling pathway during inflammation. Frontiers in immunology, 378.

Maekawa, R., Seino, Y., Ogata, H., Murase, M., Iida, A., Hosokawa, K., … Hamada, Y. (2017). Chronic high-sucrose diet increases fibroblast growth factor 21 production and energy expenditure in mice. The Journal of nutritional biochemistry, 49, 71–79.

Mente, A., Razak, F., Blankenberg, S., Vuksan, V., Davis, A. D., Miller, R., … Yusuf, S. (2010). Ethnic variation in adiponectin and leptin levels and their association with adiposity and insulin resistance. Diabetes care, 33(7), 1629–1634.

Moore, K. J., & Tabas, I. (2011). Macrophages in the pathogenesis of atherosclerosis. Cell, 145(3), 341–355.

Palomino, D. C., & Marti, L. C. (2015). Chemokines and immunity. Einstein (Sao Paulo), 13(3), 469–473. doi:10.1590/S1679-45082015RB3438

Schutte, A. (2006). van VD, van Rooyen JM, Huisman HW, Schutte R, Malan L, et al. Inflammation, obesity and cardiovascular function in African and Caucasian women from South Africa: the POWIRS study. J Hum Hypertens, 20(11), 850–859.

Schutte, A. E., Myburgh, A., Olsen, M. H., Eugen-Olsen, J., & Schutte, R. (2012). Exploring soluble urokinase plasminogen activator receptor and its relationship with arterial stiffness in a bi-ethnic population: the SAfrEIC-study. Thrombosis research, 130(2), 273–277.

Seyedsadjadi, N., & Grant, R. (2021). The potential benefit of monitoring oxidative stress and inflammation in the prevention of non-communicable diseases (NCDs). Antioxidants, 10(1), 15.

Shen, Q., Bjorkesten, J., Galli, J., Ekman, D., Broberg, J., Nordberg, N., … Landegren, U. (2018). Strong impact on plasma protein profiles by precentrifugation delay but not by repeated freeze-thaw cycles, as analyzed using multiplex proximity extension assays. Clin Chem Lab Med, 56(4), 582–594. doi:10.1515/cclm-2017-0648

Temba, G. S., Kullaya, V., Pecht, T., Mmbaga, B. T., Aschenbrenner, A. C., Ulas, T., … Kumar, V. (2021). Urban living in healthy Tanzanians is associated with an inflammatory status driven by dietary and metabolic changes. Nature immunology, 22(3), 287–300.

Temba, G. S., Vadaq, N., Wan, J., Kullaya, V., Huskens, D., Pecht, T., … Broeders, W. (2022). Differences in thrombin and plasmin generation potential between East African and Western European adults: the role of genetic and non-genetic factors. Journal of Thrombosis and Haemostasis.

Ter Horst, R., Jaeger, M., Smeekens, S. P., Oosting, M., Swertz, M. A., Li, Y., … Netea, M. G. (2016). Host and Environmental Factors Influencing Individual Human Cytokine Responses. Cell, 167(4), 1111–1124 e1113. doi:10.1016/j.cell.2016.10.018

Unwin, N., James, P., McLarty, D., Machybia, H., Nkulila, P., Tamin, B., … McNally, R. (2010). Rural to urban migration and changes in cardiovascular risk factors in Tanzania: a prospective cohort study. BMC Public Health, 10(1), 1–12.

Wang, T., Cook, I., & Leyh, T. S. (2016). Design and interpretation of human sulfotransferase 1A1 assays. Drug Metabolism and Disposition, 44(4), 481–484.

Yan, J., Nie, Y., Cao, J., Luo, M., Yan, M., Chen, Z., & He, B. (2021). The roles and pharmacological effects of FGF21 in preventing aging-associated metabolic diseases. Frontiers in cardiovascular medicine, 8, 221.

Zhang, J., Rojas, S., Singh, S., Musich, P. R., Gutierrez, M., Yao, Z., … Jiang, Y. (2021). Wnt2 Contributes to the Development of Atherosclerosis. Frontiers in cardiovascular medicine, 8.

Zhang, X., Yeung, D. C., Karpisek, M., Stejskal, D., Zhou, Z.-G., Liu, F., … Lam, K. S. (2008). Serum FGF21 levels are increased in obesity and are independently associated with the metabolic syndrome in humans. Diabetes, 57(5), 1246–1253.

